# Heterogeneous responses to embryonic critical period perturbations within the *Drosophila* larval locomotor circuit

**DOI:** 10.1101/2024.09.14.613036

**Authors:** Niklas Krick, Jacob Davies, Bramwell Coulson, Daniel Sobrido-Cameán, Michael Miller, Matthew C. W. Oswald, Aref A. Zarin, Richard A. Baines, Matthias Landgraf

**Author notes:** joint senior and corresponding authors. joint first authors (Niklas Krick, Jacob Davies, Bramwell Coulson).

## Abstract

As developing neural circuits become functional, they undergo a phase of heightened plasticity that facilitates network tuning in response to intrinsic and/or extrinsic stimuli. These developmental windows are termed critical periods (CPs), because perturbations during, but not outside, the CP can lead to lasting changes, such as the formation of sub-optimal or unstable networks. How separate, but connected elements, within a network might respond differently to a CP perturbation is not well understood. To study this, we used the locomotor network of the *Drosophila* larva as a model and heat stress as a CP stimulus that has ecological relevance. When embryos experienced heat stress the subsequent development of their locomotor network is changed, creating larvae with reduced crawling speed and decreased network stability. Developing body wall muscles and central neurons are sensitive to heat stress perturbations during distinct, consecutive phases of embryogenesis. Within the CNS, transient embryonic CP perturbation leads to increased synaptic drive from premotor interneurons to motoneurons, which in turn adopt reduced excitability. In contrast, the peripheral neuromuscular junction maintains normal synaptic transmission, despite significant structural changes. Thus, connected elements respond differentially to a CP perturbation, suggesting a sequence, or hierarchy, of network adjustment during the CP.

## Introduction

During nervous system development, critical periods (CPs) are transient windows of heightened plasticity, which are fundamental to the emergence of appropriate network function. Because of this, networks are particularly vulnerable to activity perturbations during a CP, which can result in lasting structural and/or functional changes, while comparable perturbations outside the CP are less impactful (Coulson et al., 2022; Reh et al., 2020). The functions and concepts of CPs have traditionally been investigated in the visual system of mammals (Hensch, 2005; Hensch and Quinlan, 2018; Hubel and Wiesel, 1970) and imprinting of birds (Hess, 1975; Horn, 2004). More recently, CPs have also been identified and investigated across a range of species and neuronal ensembles, from networks underlying the development of speech (Harrison et al., 2005) to motor behaviour of rats (Soiza-Reilly and Azcurra, 2009), fish (Hageter et al., 2023) and insects (Dombrovski and Condron, 2021; Giachello and Baines, 2015). The presence of a CP in developing networks is therefore increasingly considered a universal phenomenon. The precise changes within networks and associated mechanisms, as they occur during normal development, as well as those resulting from perturbations during a CP, are not well understood. This is in part due to the complexity of many of the experimental model systems, notably mammalian cortical systems that have been the predominant model for studying CPs.

To overcome this complexity, we have utilized the comparatively simpler nervous system of the fruit fly larva, *Drosophila melanogaster*, with which to study CPs. Though an insect, *Drosophila* displays CPs with features that suggest commonalities to mammalian CPs (for review see: Coulson et al., 2022). Currently, the best characterised CP during *Drosophila* development is one that occurs during embryogenesis, linked to the emergence of normal locomotor network output (Carreira-Rosario et al., 2021; Corke et al., 2025; Coulson et al., 2022; Crisp et al., 2008, 2011; Ackerman et al., 2021; Fushiki et al., 2013; Giachello et al., 2021; Giachello and Baines, 2015; Hunter et al., 2024; Sobrido-Cameán et al., 2025; Williamson et al., 2021; Zeng et al., 2021). This experimental system benefits from the comprehensively annotated connectome of the *Drosophila* larval nervous system (Giachello et al., 2020; Heckman and Doe, 2022; Valdes-Aleman et al., 2021; Winding et al., 2023) as well as an unparalleled set of genetic tools for manipulating identified neurons with both spatial and temporal precision (Simpson, 2009). Combined with imaging and electrophysiological techniques, this model system enables investigation of cellular responses to CP manipulations at the level of identified cells and synapses (Carreira-Rosario et al., 2021; Coulson et al., 2022; Crisp et al., 2008, 2011; Fushiki et al., 2013; Giachello et al., 2021; Giachello and Baines, 2015; Hunter et al., 2024; Sobrido-Cameán et al., 2025; Williamson et al., 2021; Zeng et al., 2021).

Neuronal activity manipulations during the CP of *Drosophila* embryos have been shown to cause decreased network stability. This manifests in significantly longer times required for larvae to recover from a strong activity challenge, such as electric shock-induced seizures (Coulson et al., 2022; Giachello et al., 2021; Giachello and Baines, 2015; Hunter et al., 2024, 2021). The premise that balanced network activity during the CP is necessary for stable, resilient networks to form has been explicitly supported by findings from our recent studies (Giachello et al., 2021; Giachello and Baines, 2015; Hunter et al., 2024). By extension, rectifying activity manipulations during identified CPs could have therapeutic potential (Nardou et al., 2023).

While experimental perturbations of neuronal activity during development have been a useful methodological approach for studying CPs as a phenomenon and the associated biological mechanisms, we tested if temperature change, in the form of 32°C heat stress, might be an alternative perturbation stimulus, with ecological relevance. For example, most animals are not warm-blooded and in poikilothermic animals, such as *Drosophila,* changes in ambient temperature, daily as well as seasonal, lead to changes in behavioural activity, likely caused by changes in neuronal and network activity (Lee and Montell, 2013; Powsner, 1935). Studies from the crustacean stomatogastric nervous system showed that these minimal networks produce a characteristic, rhythmic firing pattern, which is robustly maintained across a range of external temperatures. Importantly, neuronal network properties were adapted relative to the long-term temperatures that crabs had experienced, resulting in significantly different upper “crash temperatures” at which networks ceased to function (Marder, 2023; Marder and Rue, 2021). In *Drosophila,* the temperature experienced by late embryos determines larval locomotor output, including crawling speed and social feeding behaviour (Williamson et al., 2021). Similarly, studies on the *Drosophila* adult visual and olfactory systems showed that neuronal wiring properties and associated behaviours of adults are determined by the temperature experienced during a second CP of *Drosophila,* which takes place during pupariation (Chodankar et al., 2020; Kiral et al., 2021; Züfle et al., 2025), a phase of extensive network remodelling (Baumann et al., 2024; Leier et al., 2025; Lowe et al., 2023; Nelson et al., 2024).

In this study, we used heat stress of 32°C as an ecologically relevant stimulus with which to probe the embryonic CP. To gain insights at the level of single, identified connected cells, we focused on the neuromuscular system of the *Drosophila* larva, and found that the components of this network respond differently to heat stress experienced during embryonic development. Specifically, we find that elevated temperature increases neuronal activity and, when experienced during the locomotor CP, causes long term reduced network stability, as seen for neuronal activity manipulations (Giachello et al., 2021; Giachello and Baines, 2015; Hunter et al., 2024). At the neuromuscular junction (NMJ), embryonic heat stress experience causes presynaptic terminal overgrowth and changes in postsynaptic neurotransmitter receptor composition, though synaptic transmission remains normal. In contrast, within the central circuitry, we find that embryonic heat stress leads to enhanced synaptic drive from premotor interneurons and, potentially as a response, reduced excitability of motoneurons. Overall, this study presents a first comprehensive analysis of how several interconnected elements of a defined locomotor network develop in response to CP perturbation by heat stress, an ecologically relevant stimulus.

## Results

### Temperature manipulation during the locomotor CP leads to an unstable crawling network

We had previously identified a CP of nervous system development in late embryogenesis, from 17 – 19 hrs after egg laying (AEL), equivalent to 80-90% of embryonic development (Giachello and Baines, 2015). Perturbation of network activity in late embryogenesis, during this CP, pharmacologically or optogenetically, leads to the formation of an unstable, seizure prone network, as revealed by acute activity challenges. For example, an electric shock administered to late larvae, which develop from CP-manipulated embryos, causes seizures in those larvae, with recovery times that are significantly longer compared to unmanipulated controls. These pharmacological and optogenetic large-scale activity manipulations, while useful to study the embryonic CP, are artificial. We therefore asked to what extent changes in environmental conditions, that embryos might normally encounter, could also impact on network development. As poikilothermic animals, *Drosophila* development strongly depends on ambient temperature (Powsner, 1935). Several studies have shown that temperature experiences during development alter neuronal wiring as well as behaviours in larvae and in adult flies (Kiral et al., 2021; Williamson et al., 2021; Züfle et al., 2025). Increases in ambient temperature can lead to increased activity in the locomotor network (Oswald et al., 2018; Sigrist et al., 2003; Zhong and Wu, 2004). We verified this by functional imaging in isolated CNS, measuring spontaneous calcium transients from motoneurons, via GCaMP8f selectively expressed in aCC motoneurons using *R94G06-GAL4*. Stepwise increases in temperature led to increases in the frequency of rhythmic bursts. In contrast, CNS kept at the control temperature of 25°C for the duration of functional imaging (600 seconds) tended to show a decrease in frequency over time (Fig. 1A). These experiments suggest that increases in ambient temperature, above 25°C, lead to concomitant increases in locomotor network activity.

**Figure 1:**
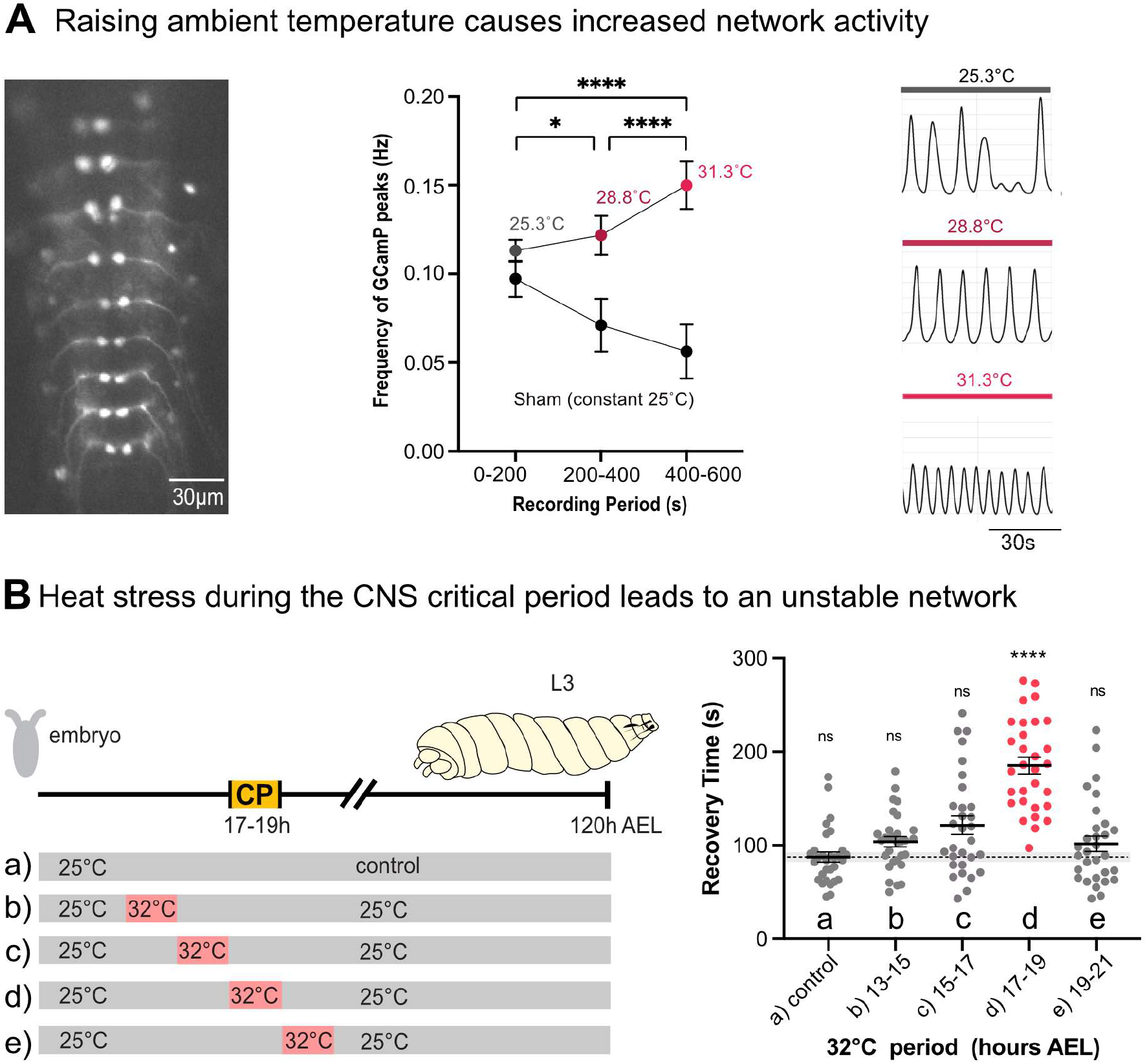
Transient heat stress during the critical period leads to an unstable network. **A)** Acute increase of ambient temperature leads to increased network activity, as measured with the fluorescent calcium indicator GCaMP8f expressed in segmentally repeated aCC motoneurons in isolated central nervous systems; while constant 25°C leads to reduced activity over time. Left: micrograph of an isolated ventral nerve cord with segmentally repeated aCC motoneurons expressing GCaMP8f. Centre: traces of GCaMP fluorescence peaks measured at different temperatures. Right: quantification of fluorescence peak frequency changes over time (control/sham -bottom trace) and in response to temperature increase (top trace). **B)** Heat stress, when experienced during the embryonic CP of locomotor network development, leads to reduced network stability. Left: Schematic representation of embryonic and larval development and the paradigm of temperature manipulation during embryogenesis. Embryos were kept at the control temperature of 25°C, either continuously (control) or moved to 32°C for 2 hours, either before (13-15 h, 15-17 h AEL), during (17-19 h AEL) or after (19-21 h AEL) the locomotor network CP, then returned to 25°C until the late larval stage, when subjected to seizure induction by electroshock. Right: quantification of seizure recovery times. n=30, data are shown with mean ± SEM, ANOVA, ****p < 0.00001, ‘ns’ indicates statistical non-significance.

Next, we tested the effect that a transient experience of 7°C above the evolved temperature preference of 25°C (Ashburner et al., 1983), namely 32°C, might have on nervous system development. We chose 32°C as this is known to be experienced by *Drosophila melanogaster* (Williamson et al., 2021). We transiently exposed embryos for two hours to 32°C heat stress either before, during, or after the previously characterised embryonic CP for locomotor network development (17 – 19 hrs AEL) (Giachello and Baines, 2015), then returned them to the control temperature of 25°C. After four days, at the end of larval development, we measured as a proxy for network stability the speed of recovery from electroshock-induced seizures. Seizure recovery was unchanged, at control levels, following transient heat stress before or after the embryonic CP. In contrast, animals that had experienced 32°C heat stress during the embryonic CP (i.e. between 17 – 19 hrs) had significantly prolonged recovery times, indicative of reduced network stability (Fig. 1B). This shows that when *Drosophila* embryos experience a 32°C heat stress during the embryonic CP of the locomotor network it results in altered developmental outcomes, notably the formation of sub-optimal, less stable networks, similar to the effect of network-wide activity perturbations during this developmental window (Hunter et al., 2024; Giachello et al., 2021; Giachello and Baines, 2015).

To investigate structural and functional changes that result from a CP heat stress perturbation we first focused on the well characterised, and easy to access, peripheral neuromuscular junction (NMJ). Specifically, we focused on the NMJ of dorsal acute muscle 1 (DA1), which is innervated by the aCC motoneuron, a cell that has been the focus of our previous studies (Fig. 2A) (Baines et al., 2001, 1999; Fujioka et al., 2003; Hunter et al., 2024; Landgraf et al., 2003; Giachello et al., 2021; Giachello and Baines, 2015 Sobrido-Cameán et al., 2025). Because temperature manipulations outside the CP did not lead to an unstable network (Fig. 1A), we adopted a simplified experimental paradigm of putting embryos from a 6h egg collection to 32°C for 24h. This produces outcomes comparable to heat stress exposure centred around the CP, while ensuring that all embryos go through their CP during heat stress irrespective of minor variations in developmental timing or due to retention of fertilised eggs. We find that heat stress during embryogenesis leads to a significant overgrowth of the aCC motoneuron terminal (composed of type Ib boutons) by the late larval stage. This increase in the number of boutons is accompanied by a corresponding increase in active zones, while overall active zone density is not altered (Fig. 2B). On the postsynaptic side, embryonic heat stress causes an altered receptor composition: a selective reduction of ion channels containing the larger conductance GluRIIA subunit. Across the aCC NMJ, this leads to a reduced ratio of the larger conductance GluRIIA *versus* the lower conductance GluRIIB subunit-containing glutamate receptors (Fig. 2C).

**Figure 2:**
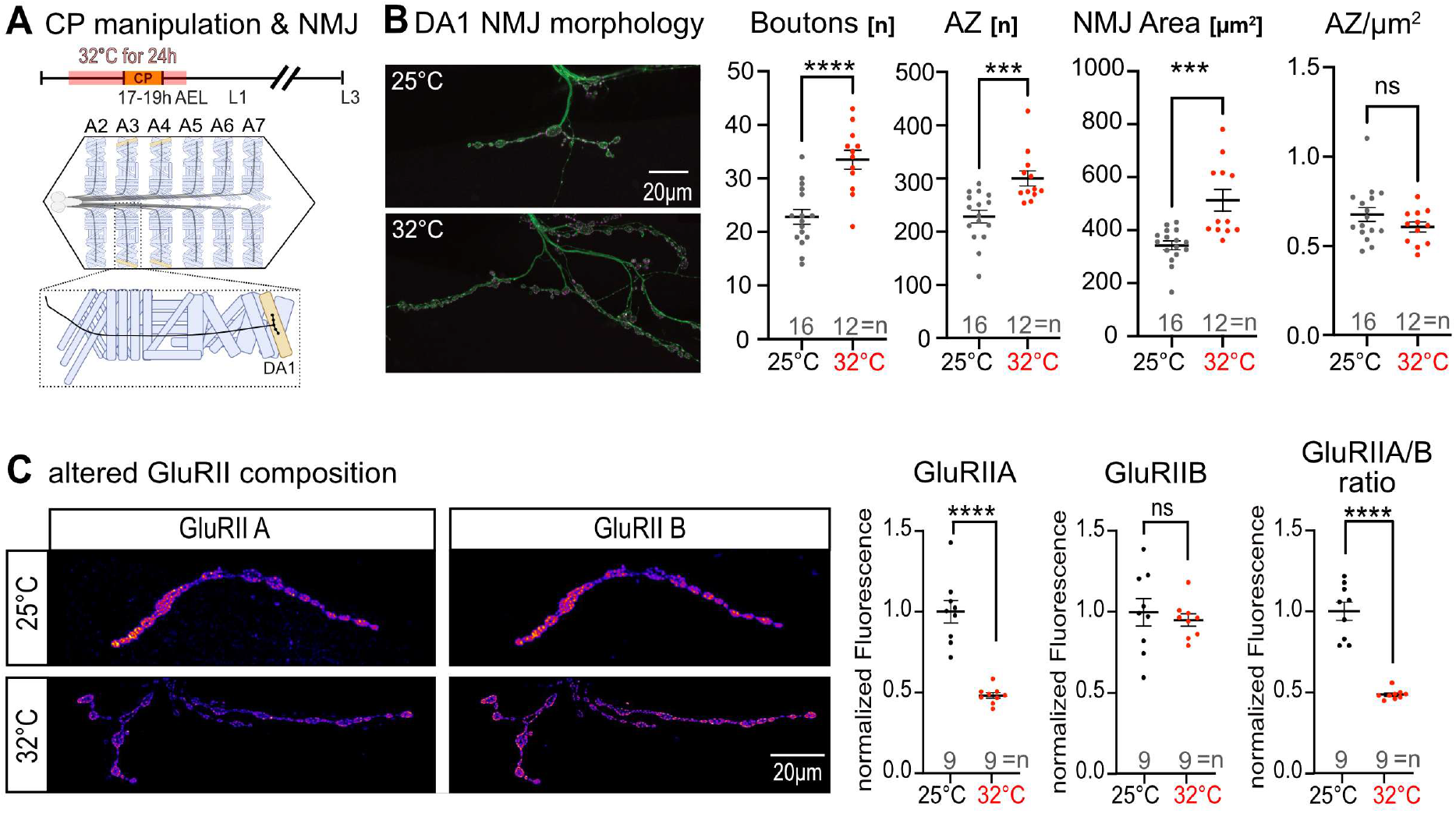
Morphological changes at the neuromuscular junction after critical period perturbation. **A)** Schematic of temperature manipulation during embryogenesis and location of dorsal acute muscle 1 (DA1) within a fillet dissected larva. Embryos collected over a 6h period were incubated at 32°C for 24h to guarantee heat stress exposure during the locomotor network CP. **B)** This results in a change of subsequent NMJ development, which manifests as a significant overgrowth of aCC presynaptic terminals (green, visualised with anti-HRP-AlexaFluor488). The increase in bouton number and NMJ area is paralleled by concomitant increases in active zones (magenta), resulting in a normal active zone density. **C)** Postsynaptically, embryonic heat stress causes in a significant decrease of glutamate receptors containing the larger conductance GluRIIA subunit, while those with the GluRIIB subunit remain unaffected. Thus, the postsynaptic GluRII receptor composition is significantly changed following a transient embryonic heat stress, as reflected in the ratio of receptors containing GluRIIA *vs* GluRIIB. Data are shown with mean ± SEM, unpaired t-test, ***p < 0.0001, ****p < 0.00001, ‘ns’ indicates statistical non-significance.

In recognition of the potential caveat that prolonged heat stress during most of embryogenesis could lead to compound effects, we additionally carried out 32°C heat stress exposure of embryos during shorter time windows, followed by rearing at the control temperature of 25°C. We then measured postsynaptic GluRIIA *vs* GluRIIB levels at the aCC NMJ on muscle DA1 of late wandering third instar larvae. We found that the GluRIIA reductions caused by prolonged heat stress throughout most of embryogenesis is recapitulated by a shorter, 3h exposure to 32°C, though during a different developmental window, from 13-16 hrs AEL. Heat stress before or after this 13-16 hrs AEL developmental window had no effect on the composition of the postsynaptic glutamate receptors (Fig. 3). GluRIIB subunit levels, in contrast, remained unaffected by embryonic exposure to 32°C heat stress. The period of 13-16 hrs AEL is the developmental window during which the embryonic body wall muscles become electrically active and start to exhibit myogenic contractions (Broadie and Bate, 1993; Crisp et al., 2008). Thus, the maturation of the embryonic body wall muscles precedes the CP of the central locomotor circuitry (17-19 hrs AEL), when central neurons develop their electrical properties (Baines and Bate, 1998). To our surprise, we found that NMJ size is impacted by heat stress during an even earlier window of muscle development (9-12 hrs AEL), during which myoblast fusion occurs (Bate et al., 1993) (Fig. 3). Overall, we found that transient embryonic heat stress of 32°C leads to reproducible changes in NMJ development during subsequent larval stages. Our observations show that postsynaptic glutamate receptor composition is specified during the developmental window that is characterised by electrical maturation of the body wall muscles, preceding by several hours the equivalent maturation of the innervating motoneurons.

**Figure 3:**
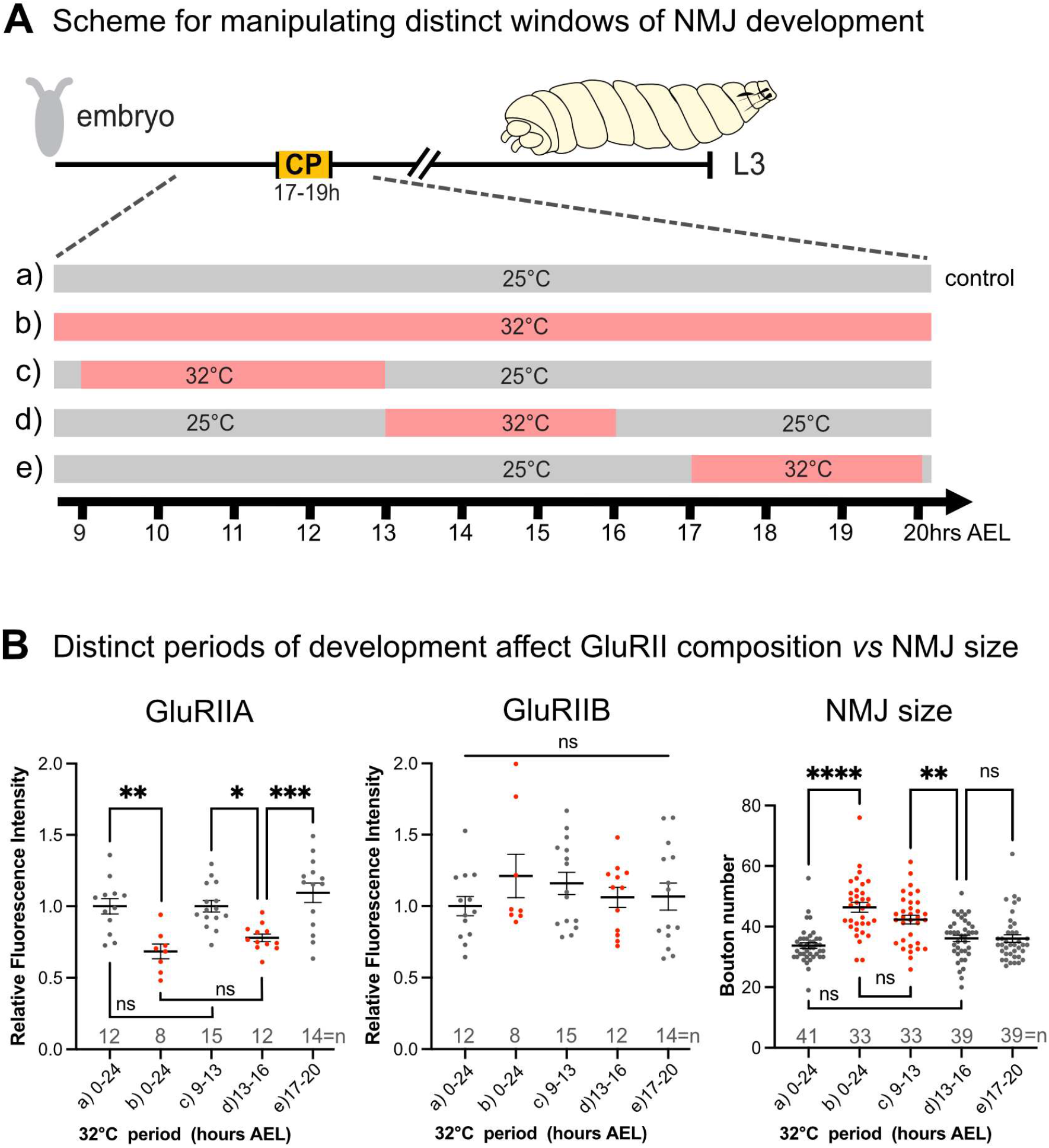
Heat stress pertubations during distinct periods of development differentially affect postsynaptic GluRII receptor composition *vs* NMJ size. **A)** Schematic of temperature manipulations during embryogenesis. Embryos collected over one hour were incubated at 32°C for different phases of embryogenesis. **B)** Resultant changes to the development of the aCC NMJ on muscle DA1 were quantified at the late wandering third instar larval stage. Reduced expression of the GluRIIA subunit occurred following heat stress experienced during 13-16 hours AEL, comparable to GluRIIA changes caused by heat stress during all of embryogenesis, but not in response to heat stress experienced before or after this 3-hour window. GluRIIB levels are not affected by heat stress during any period or all of embryogenesis. NMJ size, in contrast, is affected by heat stress experienced during earlier phases of muscle development, from 9-13 hours AEL. Data are shown with mean ± SEM, ANOVA, *p < 0.01, **p < 0.001, ***p < 0.0001, ****p < 0.00001, ‘ns’ indicates statistical non-significance. Sample sizes: for GluRIIA and GluRIIB measurements, each data point represents a unique specimen. NMJ size was measured in the same specimens for up to four muscle DA1 NMJs in abdominal segments 3-5 sampled per larva.

### Despite morphological changes, larval NMJ physiology is not altered

In view of the prominent changes in NMJ growth, morphology and postsynaptic glutamate receptor composition that result when heat stress (32°C) is experienced during embryonic development, we next assessed NMJ synaptic physiology. Because the NMJ is a composite structure that develops through interactions between the postsynaptic body wall muscle with presynaptic terminals from motoneurons, whose cell bodies are located within the CNS, we proceeded using an experimental paradigm of prolonged heat stress (32°C) during most of embryogenesis, so as to include CPs of both body wall muscles and the central locomotor circuitry. We used the two-electrode voltage clamp (TEVC) technique to assess muscle synaptic physiology, initially measuring single pulse transmission across a gradient of external calcium concentrations. Despite the marked structural changes in heat stress manipulated animals, single pulse transmission at the NMJ was not significantly different between embryonically manipulated (32°C experience during embryogenesis) and controls that had been reared at 25°C throughout (Fig. 4 A&B). Quantification of the number of release sites utilised per action potential and quantal size (Jusyte et al., 2023) also showed no statistically significant difference between manipulated and control specimens (Fig. 4C). We quantified the size of the readily releasable pool (RRP) of synaptic vesicles by recording responses to a high frequency train of action potentials (Fig. 4D), then plotting the cumulative amplitude of responses to the first 60 pulses and linearly back extrapolating through the last 20 pulses to y=0 (Fig. 4E). This also did not show a significant difference between manipulated and control animals (Fig. 4F). Though we noticed that during the 60Hz pulse train, NMJs in embryonic heat stress-manipulated specimens exhibited an increased facilitation during the first 10 pulses, indicating a slight reduction in release probability of the presynaptic terminal (Fig. 4D). Last, we tested if the NMJ remained able to adjust homeostatically to acute changes in synaptic transmission. We utilized the well-established method of acutely blocking glutamate receptors with Philanthotoxin (PhTx), a GluRIIA blocker, to induce presynaptic homeostatic potentiation (Frank et al., 2006; Genç and Davis, 2019; Müller and Davis, 2012). Acute GluRIIA blockade by a 10 min incubation with PhTx caused a significant reduction mEPSC amplitude while evoked EPSC amplitude remained unchanged in both controls and heat stress-manipulated specimens. Calculating quantal content shows that in response to acute GluRIIA blockade, both control and manipulated animals are indeed able to induce presynaptic homeostatic potentiation by increasing quantal content (Fig. 4G-Giii). While establishing baseline conditions, we found mEPSC amplitudes and quantal content to be comparable between controls and experimentals Fig. 4G-Giii). This was contrary to expectation considering the overall reduction in the GluRIIA/B ratio that results from an embryonic heat stress manipulation. Overall, we conclude that despite the significant structural changes that emerge by the late larval stage following an embryonic 32°C heat stress experience, synaptic physiology at the late larval NMJ remains normal. Moreover, when considering the unstable network phenotype, as manifest by increased recovery times from electroshock-induced seizures, our finding that synaptic transmission at the NMJ remained normal suggests that the network instability likely originates from changes within the central locomotor circuitry.

**Figure 4:**
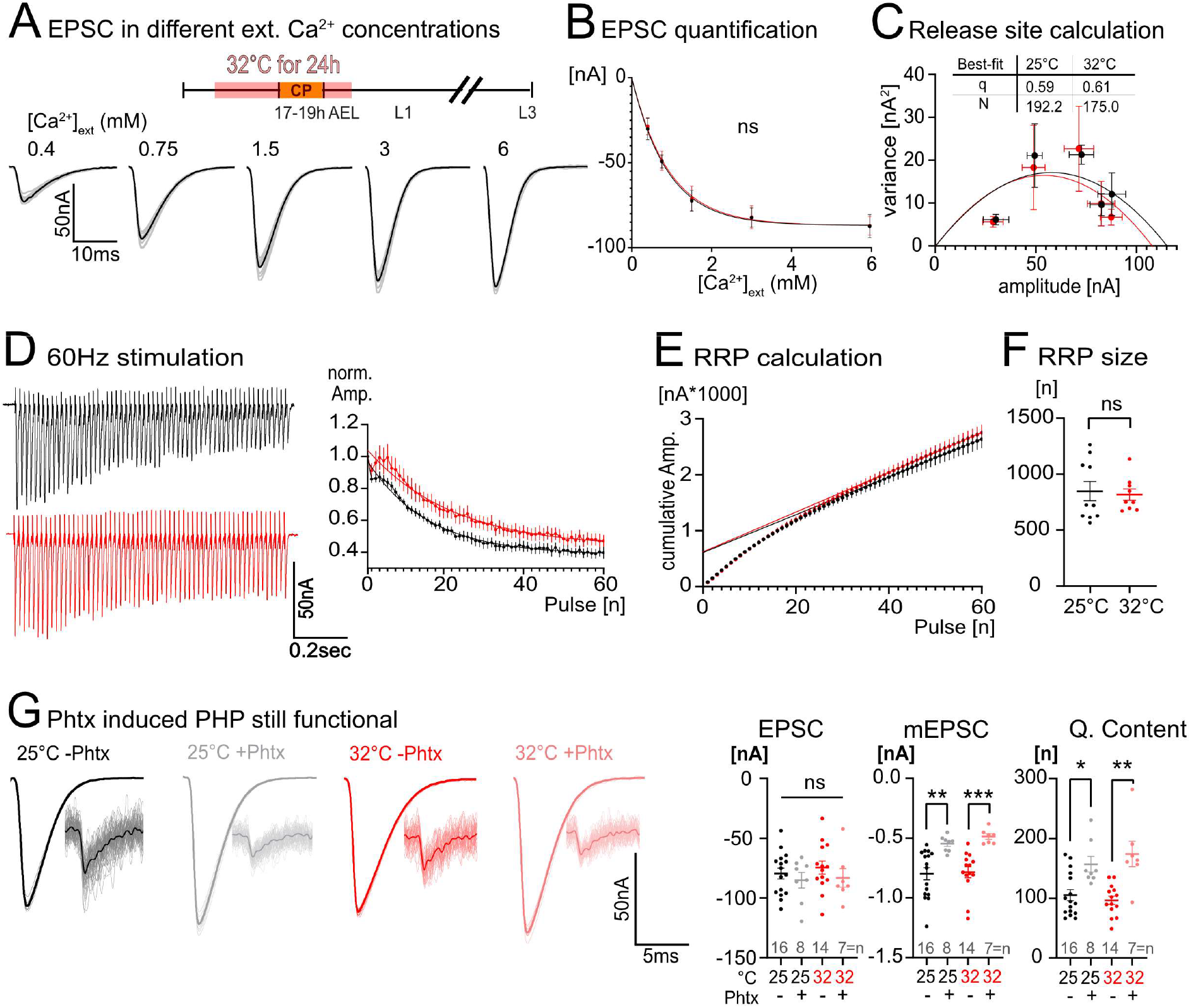
Physiological properties of the late larval NMJ following an embryonic 32°C perturbation. **A)** Representative traces of control two electrode voltage clamp recordings across a gradient of external calcium concentrations. **B)** Postsynaptic current amplitude is not altered between control and embryonic heat stress-manipulated animals at any given external calcium concentration (n=5). **C)** Plotting postsynaptic current amplitude against its variance to calculate quantal size (q) and the number of release sites being used (N), also did not show a significant difference between control and embryonic heat stress-manipulated animals (n=5). **D)** To measure readily releasable pool size, a 60Hz stimulus train was applied for 1s where we could observe a slight increase in facilitation over the first 10 pulses in specimens that in their embryonic stage had experienced heat stress of 32°C (n=9) as compared to controls (n=11). **E)** Cumulative amplitude was plotted over time and a linear regression was back extrapolated through the last 20 pulses (same data as for E). **F)** The y intercept was divided by the Mini size of the same trace to calculate the readily releasable pool (RRP) size. **G)** NMJs in heat stress-manipulated specimens have normal EPSC and mEPSC amplitudes and quantal content. Acute application of the GluRIIA blocker PhTx reliably induced presynaptic homeostatic potentiation in controls and 32°C manipulated animals. Data in F & G are shown with mean ± SEM, ANOVA, *p < 0.01, **p < 0.001, ***p < 0.0001, ‘ns’ indicates statistical non-significance.

### Motoneurons display reduced excitability and receive increased synaptic input following embryonic perturbation by heat stress

We noted that following embryonic exposure to 32°C heat stress, motor axons in late third instar larvae required an increased stimulation voltage to reliably induce action potentials. This hinted at motoneurons in these larvae having acquired a state of reduced excitability. To directly assay motoneuron excitability, we used whole cell patch clamp to measure action potential firing of aCC, evoked by injection of current steps (4pA). These measures were made in the presence of mecamylamine (200 µM in external saline) to block endogenous cholinergic excitatory synaptic drive. We found that in late third instar larvae, the aCC motoneuron had significantly decreased excitability following an embryonic heat stress experience of 32°C (Fig. 5A). These results are comparable to, and validate, results obtained after optogenetic locomotor network activity manipulation during the embryonic CP of the locomotor network (17-19 hrs AEL) (Giachello and Baines, 2015). In previous studies, we had shown with recordings from aCC that its synaptic drive is enhanced following transient network activity manipulations during the embryonic CP, using optogenetic or pharmacological means to induce network over-excitation or genetic persistent overactivation of neurons, as in *para^bss^* seizure mutants (Giachello and Baines, 2015; Hunter et al., 2024). This manifests as an increase in the duration (though not amplitude) of endogenous excitatory cholinergic synaptic currents, termed spontaneous rhythmic currents (SRCs) (Giachello et al., 2019; Giachello and Baines, 2015). We asked if heat stress experienced during embryonic development led to similar changes. Indeed, in late larvae that had experienced heat stress during embryonic development, SRCs recorded from aCC motoneurons were significantly increased in duration, though with amplitude unchanged (Fig. 5Bi), i.e. phenocopying the lasting effects of network activity manipulations during the embryonic CP. Note these SRC recordings were undertaken in the absence of mecamylamine and suggest an increase in premotor network activity.

**Figure 5:**
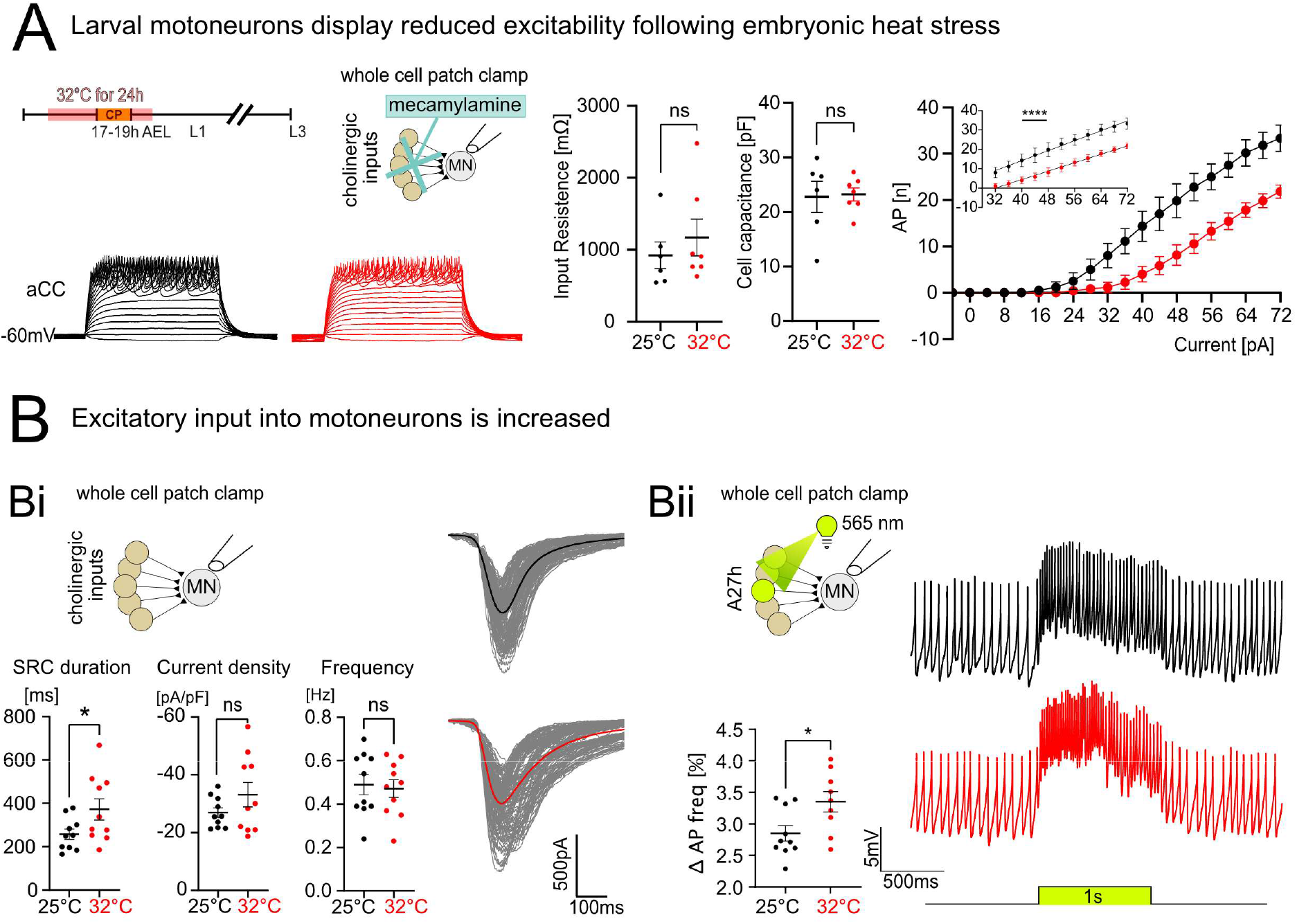
Larval motoneurons display reduced excitability and increased synaptic input after an embryonic 32°C heat stress experience. **A)** Embryos were subjected to 32°C heat stress, then reared to the late larval stage under control conditions of 25°C. Whole cell patch clamp recordings show action potentials generated by aCC motoneurons isolated from excitatory synaptic input by mecamylamine addition in response to current injection. Inset shows linear regression centred around 32-72 pA current injections, showing significant reduction in excitability in heat stress manipulated specimens relative to controls. **B)** Whole cell patch clamp recordings from the aCC motoneuron in larvae: **Bi)** reveal an increase in the duration of spontaneous rhythmic currents (SRCs) in larvae that had previously experienced 32°C heat stress during embryonic development. Current density and frequency of SRCs remain comparable to controls. **Bii)** Targeted optogenetic activation (using Chronos, for 1 second) of one premotor excitatory interneuron, A27h, showed A27h synaptic input has increased as a result of a previous embryonic 32°C heat stress experience. Data are shown with mean ± SEM, unpaired t-test, *p < 0.01, ****p < 0.00001, ‘ns’ indicates statistical non-significance.

To further explore changes in the premotor network, we recorded synaptic drive to aCC under whole cell patch clamp configuration. We focused on one of the most the strongly connected excitatory premotor interneurons, termed “A27h”. Because motoneurons are quiescent without synaptic drive, a small amount of current was injected into aCC to maintain a modest and consistent mean firing frequency of ∼10-20 Hz. We then assayed how this mean firing frequency changed upon optogenetic activation of the A27h premotor interneuron (Hunter et al., 2024). As might be expected, and as per our previous study (Hunter et al., 2024), acute optogenetic activation of the A27h premotor excitor resulted in increased aCC firing frequency. Significantly, this increase was greater in larvae that had been exposed to heat stress during their embryonic development, as compared to controls (Fig 5.Bii). This observation demonstrates that heat stress during embryonic development results in increased excitatory drive from central premotor interneurons, and it suggests that this might then countered by the postsynaptic motoneurons through reduction in their excitability.

### Adjustments of the pre-motor synaptic drive onto the aCC motoneuron result in normal endogenous firing patterns

If the reduction in excitability displayed by the aCC motoneuron was a homeostatic response to the increased premotor synaptic drive, then one might predict this to be a mechanism to maintain the endogenous level of motoneuron activity. To test this, we used cell-attached patch recordings from aCC motoneurons, in isolated nerve cords of late third instar larvae.

This extracellular recording mode allows quantification of endogenous action potential firing without disturbance of the intracellular composition of a neuron. As hypothesized, our recordings show that no significant differences in endogenous motoneuron activity result from an embryonic experience of heat stress, as measured by burst rate, burst duration, or action potentials per burst (Fig. 6A). Next, to test if this translates to normal muscle output, we recorded spontaneous synaptic drive from muscles in a semi-intact preparation with an intact nervous system attached. Current clamp recordings from the DA1 muscle, while supressing muscle contractions using nifedipine (50µM) (Dyson et al., 2022; Kratschmer et al., 2021), showed no significant differences in spontaneous burst frequency or duration, nor the overall time spent bursting (Fig. 6B), between controls and embryonic heat stress-manipulated specimens. Therefore, we conclude that, at least in this locomotor system, CP manipulations might act primarily on the cells of the premotor circuitry, namely excitatory premotor interneurons as tested in this study. Changes in the motoneurons could thus be seen to constitute secondary homeostatic adjustments in response to those premotor network changes.

**Figure 6:**
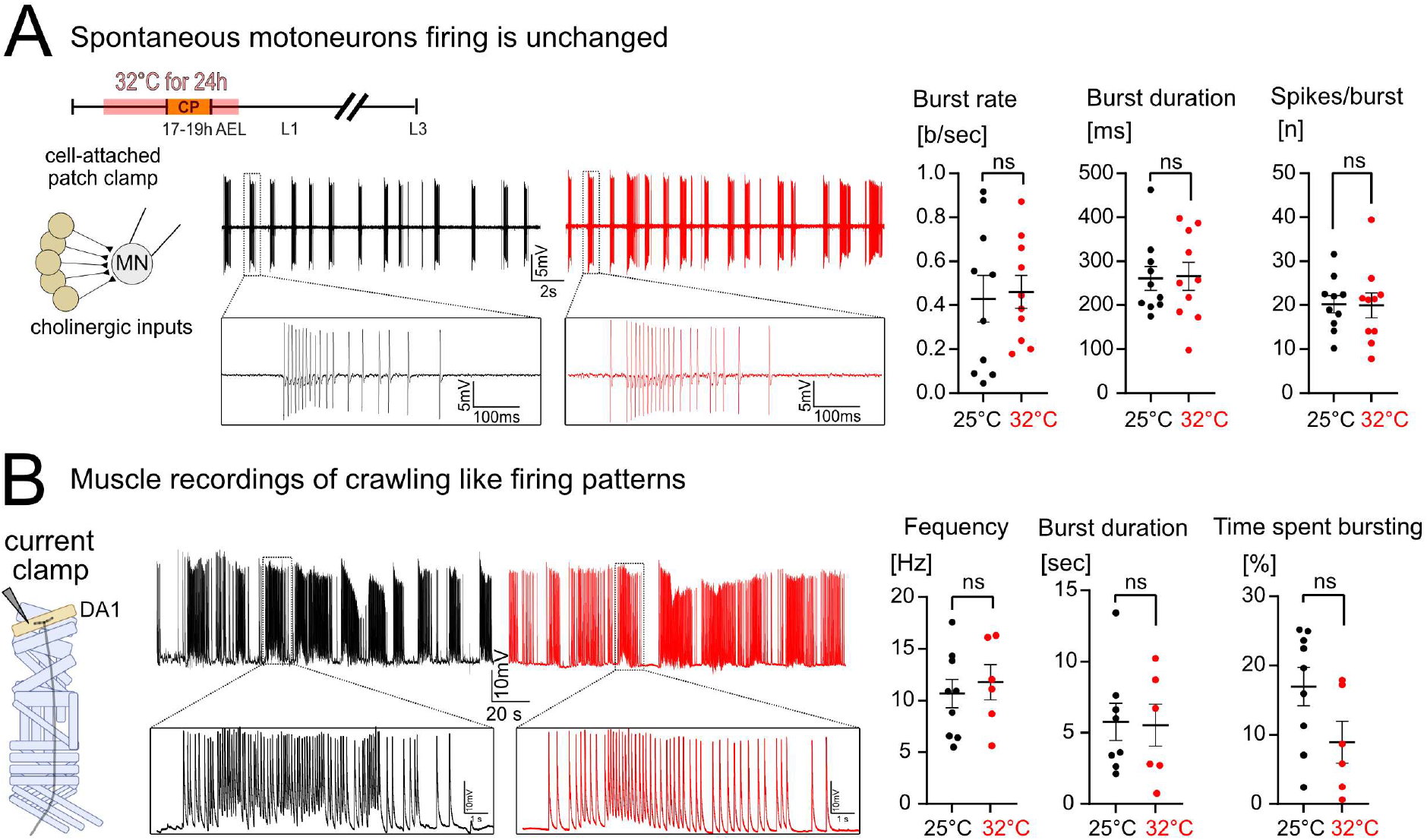
Spontaneous larval motoneuron and muscle firing properties are unchanged following embryonic heat stress. **A)** Monitoring spontaneous motoneuron firing by cell-attached patch clamp recordings from the aCC motoneuron in a late third instar larval nerve cord found no significant differences between embryonically manipulated specimens and controls, regarding burst rate per second, burst duration or number of spikes per burst (n=10). **B)** Measuring muscle output generated by motoneuron spontaneous firing with current clamp recordings from the DA1 muscle in late third instar larvae showed that larvae are able to produce normal motoneuron firing output to the muscle, irrespective of embryonic experience (n=8 for controls, n=6 for experimentals with 32°C embryonic experience). Data are shown with mean ± SEM, unpaired t-test, ‘ns’ indicates statistical non-significance.

### Propagation speed of neuronal activation waves is reduced following embryonic heat stress perturbation, mirroring reduced crawling speed

To establish how the excitatory premotor network is affected by a transient embryonic 32°C heat stress manipulation we conducted functional imaging of the segmentally repeated A27h premotor interneurons. Specifically, we targeted expression of the calcium indicator GCaMP8f to A27h premotor interneurons. Then, in isolated third instar nerve cords, we recorded the propagation of neuronal activation waves, indicative of fictive locomotion, across abdominal segments (Fig. 7A) (Streit et al., 2016; Pulver et al., 2015). These experiments show that embryonic 32°C exposure leads to a significant reduction in the speed with which these neuronal activity waves travel from posterior to anterior abdominal segments, as compared to controls, and mirrors the decrease in larval crawling speed caused by the same embryonic manipulation (Fig. 7B). Because our electrophysiological analyses showed that motoneuron firing patterns and output to the muscle remain normal after an embryonic 32°C manipulation (Fig. 4-6), these results further support the hypothesis that it is the lasting changes to the central premotor circuitry, rather than to the motoneurons, which are responsible for the behavioural changes, in crawling behaviour, and likely also for network instability.

**Figure 7:**
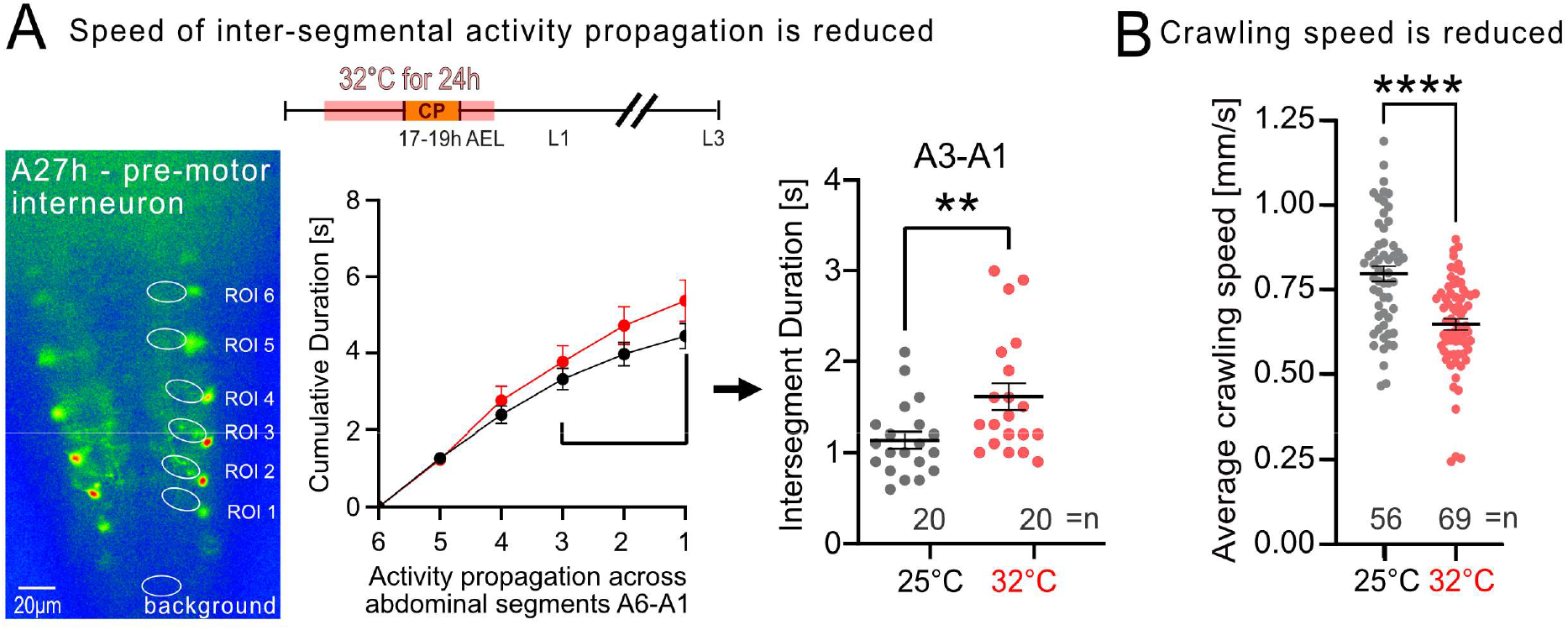
Embryonic heat stress manipulation causes reduced speed of both, activity wave propagation and larval crawling. **A)** Functional imaging of A27h premotor interneurons using GCaMP8f in isolated CNSs from late larvae that previously had experienced transient 32°C heat stress during embryonic development. Activity wave propagation across abdominal segments is slowed down, i.e. intersegment duration has increased, notably between anterior segments, A3 to A1. **B)** In intact animals, locomotor network output is similarly altered: embryonic heat stress (32°C) leads to a significant reduction in larval crawling speed, as compared to controls (25°C). Data are shown with mean ± SEM, unpaired t-test, **p < 0.001, ****p < 0.00001, ‘ns’ indicates statistical non-significance. For B) only the crawling speed of each larva was sampled up to three times, once per unique 5-minute crawling interval.

### The CP of the locomotor network determines behavioural outcomes

Having identified consecutive phases of embryonic development that represent CPs for the body wall muscles and NMJ development, followed by a CP for the central locomotor circuitry (Fig. 3), we asked which of these CPs shaped larval crawling behaviour. To this end, we subjected embryos to 32°C heat stress at different windows of their development, separating the periods that principally affects NMJ development (prior to 17 hours AEL) from the subsequent 2-hour window during which the development of the locomotor circuitry is vulnerable to neuronal activity and heat stress manipulations (17-19 hours AEL) (Giachello and Baines, 2015). Assaying crawling behaviour of third instar larvae showed that crawling speed is reduced only when the embryonic CP of the locomotor network (17-19 hours AEL) had been subjected to heat stress. This demonstrates that it is the impact on the developing central circuitry, rather than NMJ formation, which causes behavioural changes in terms of crawling speed (Fig. 8A) and of network stability, as assessed by seizure recovery times (Fig. 1).

**Figure 8:**
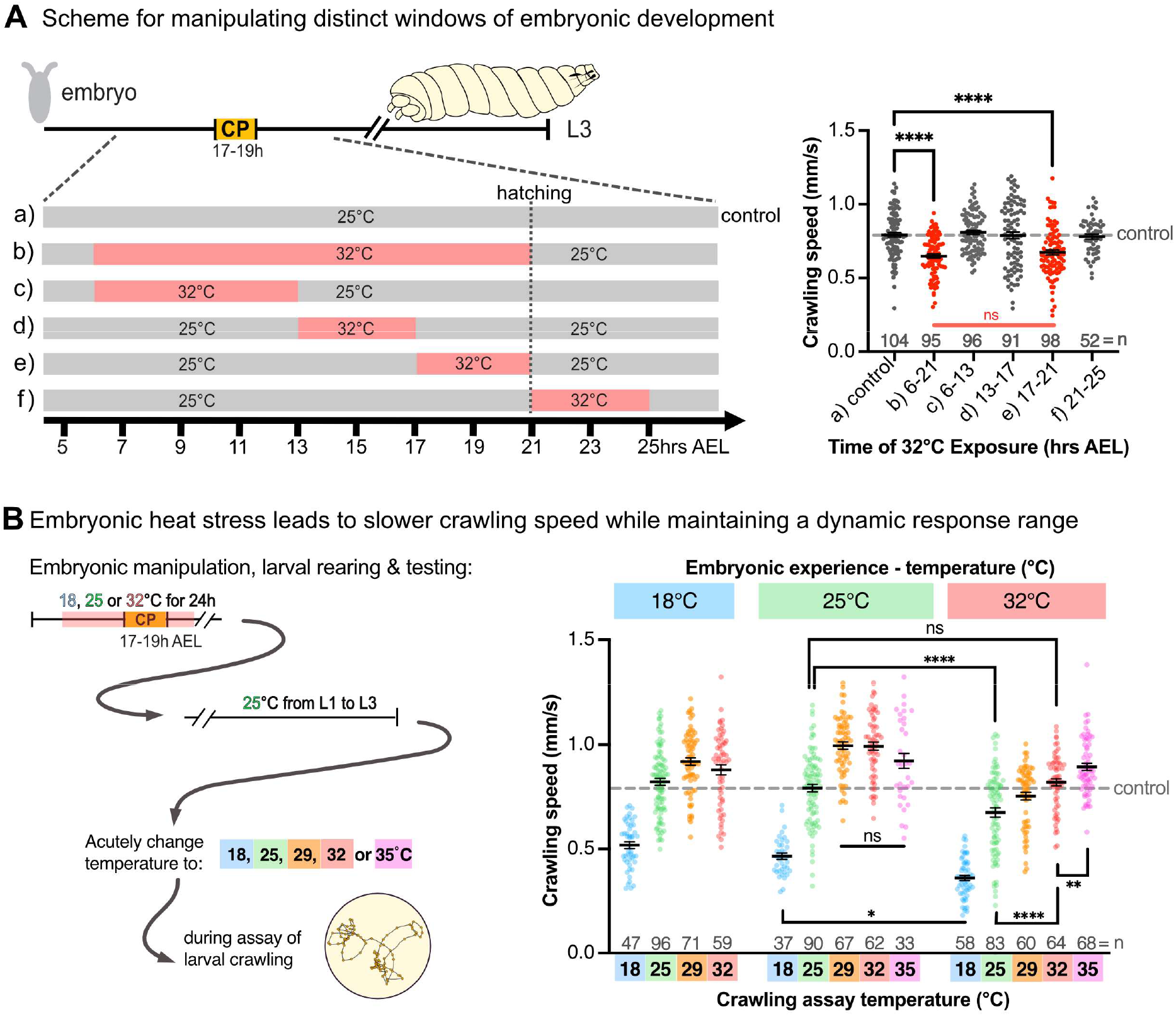
Larval crawling speed is determined by the CP of central locomotor network. **A)** Embryos were subjected to heat stress (32°C) during different windows of development. Crawling speed of resultant third instar larvae was affected only when the CP for the central locomotor network (17-19hrs after egg laying [AEL]) was manipulated. **B)** Embryonic experience of heat stress leads to larvae that retain their ability to increase crawling speed in response to acute relative increases in ambient temperature, to a higher absolute temperature (35°C) than controls (29°C). Data are shown with mean ± SEM, ANOVA, *p < 0.01, **p < 0.001, ****p < 0.00001, ‘ns’ indicates statistical non-significance. Each larva was sampled up to three times, once per unique 5-minute crawling interval.

We wondered whether larvae that emerge from heat stressed embryos crawl more slowly simply because they are no longer able to crawl at the normal speed. To test this, we subjected embryonic development to 18, 25 or 32°C and, after hatching, reared larvae at 25°C until the mid-third instar stage (72 hrs ALH). We then assayed how these larvae adjusted their crawling speed in response to an acute change in ambient temperature, when placed into an arena equilibrated to 18, 25, 29, 32 or 35°C (Fig. 8B). This clearly showed that while embryonic heat stress leads to a significant reduction in the default crawling speed, larvae retained the ability to accelerate in response to an acute increase in ambient temperature, as they normally do. Moreover, animals that experienced a higher temperature during embryogenesis retained the ability to increase their crawling speed at higher temperatures (e.g. 32 or 35°C) than those with cooler embryonic experiences, whose maximum speed plateaued at 29°C (Fig. 8B). These data suggest that while an embryonic experience of heat stress reduces the default crawling speed, it preserves a response range at higher temperatures, which might therefore potentially be adaptive to larval life at higher temperatures.

## Discussion

### An experimental model system with which to study CP biology

In this study we utilised the *Drosophila* embryo and larva as an experimental model with which to study the role of CP-regulated plasticity across interconnected cells within a defined circuit, from a premotor excitatory interneuron, to motoneuron to muscle. We have identified 32°C heat stress as a CP stimulus that has ecological relevance, as a temperature experienced in the wild (Williamson et al., 2021). Transient exposure of embryos to 32°C during the previously defined CP for the developing CNS (17-19 hours after egg laying) leads to lasting changes in subsequent nervous system development, resulting in decreased crawling speed of third instar larvae and in reduced network stability, reflected by an increased susceptibility to electroshock-induced seizures. As is characteristic for CPs, the same manipulation before or after this defined developmental window has no lasting effect on nervous system properties or animal behaviour. A well-known way by which increases in temperature impact on neurons is their activity. Using calcium imaging of motoneurons we showed that stepwise increases in ambient temperature cause acute increases of fictive rhythmic locomotor network activity in isolated nerve cords. This agrees with past studies, which have demonstrated the well-established link between temperature and neuronal activity in this same experimental system (Zhong and Wu, 2004). It is therefore not surprising that 32°C heat stress during the embryonic CP of nervous system development causes outcomes that are comparable to those created by neuronal activity manipulations (pharmacological or genetic) during this developmental window (Giachello et al., 2021; Giachello and Baines, 2015; Hunter et al., 2024).

### Different cell types respond differently to CP manipulations

A particular strength of the *Drosophila* larval motor system is the accessibility of interconnected network elements. We focused on a small part of the locomotor network: an excitatory premotor interneuron (A27h) connected to a motoneuron (aCC), and its postsynaptic body wall muscle (DA1). Following transient application of heat stress during embryonic development, we could identify that within this section of the locomotor network each of these three connected nodes responded differently. We observed increased excitatory drive from A27h interneurons to aCC motoneurons, which might reflect changes in A27h neuronal excitability and/or its synaptic transmission onto motoneurons. Potentially in response to these changes in the premotor network, aCC motoneurons displayed a decrease in excitability, which facilitated normal motoneuron firing patterns to be maintained. In contrast to these changes within the central circuitry, at the peripheral NMJ terminal, we documented normal synaptic transmission properties, despite a significant presynaptic terminal overgrowth. The latter might be a secondary adjustment to a decrease in transmitter sensitivity by the muscle, which showed reduced levels of the high conductance subunit GluRIIA.

The first observation to make is that these responses to embryonic heat stress, which manifest during subsequent larval stages, are reproducible. This suggests that there might be direct cause-effect relationships at the cellular level, which can now be investigated in detail using this experimental model. Building on these findings, we thus identified mitochondrial reactive oxygen species (ROS) and Hypoxia-inducible factor-1 alpha (HIF-1α) as heat stress-activated signals required for the long-lasting changes of cellular properties that result from embryonic CP heat stress (Sobrido-Cameán et al., 2025). An alternative, not mutually exclusive interpretation is that activity patterns in this part of the network might be robustly canalised so as to lead to reproducible outcomes. Second, the changes in neuronal excitability and synaptic transmission that we have documented are compatible with a temporal sequence, or a functional hierarchy, during network maturation. For example, following an embryonic heat stress, premotor interneurons, such as A27h, potentially part of the central pattern generator (Carreira-Rosario et al., 2021; Zeng et al., 2021), increase their collective synaptic drive to postsynaptic motoneurons, which in turn reduce their excitability (Fig. 5). Along these lines we interpret the changes in motoneuron properties that we have characterised as homeostatic responses by motoneurons to the increase in presynaptic excitatory drive from premotor interneurons. This perspective implies a hierarchy, whereby the specification of premotor interneuron properties dominates that of their downstream integrators, the motoneurons (Fig. 9). This is supported by previous findings of motoneurons flexibly adjusting their electrical properties in response to changes in presynaptic excitatory drive (Baines et al., 2001). However, the precise temporal and causal relationships of cell type-specific adjustments during the CP of this network remain to be established. In the developing mouse neocortex, sequential cell type-specific maturation windows have recently been shown. There, somatostatin-expressing inhibitory interneurons differentiate early, and promote the maturation of network activity patterns through inhibition of the later differentiating parvalbumin-expressing inhibitory interneurons (Mòdol et al., 2024).

**Figure 9:**
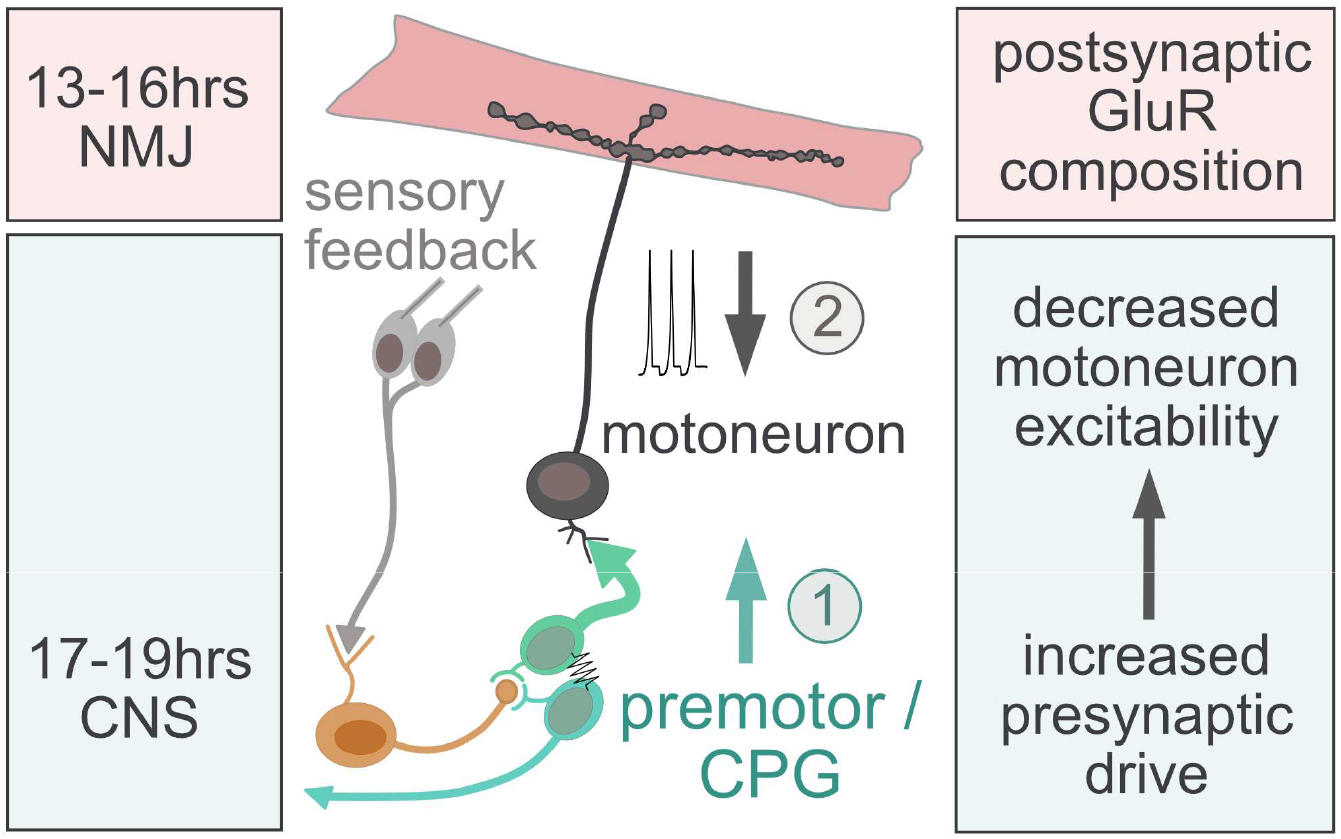
Model of distinct critical periods for body wall muscles and the central locomotor circuitry. Embryonic heat stress (32°C) leads to adjustments of postsynaptic receptor fields, when applied during an earlier developmental window of 13-16 hrs after egg laying. Subsequently, from 17-19 hrs after egg laying the central locomotor circuitry is sensitive to perturbations, leading to an increase in synaptic drive of the premotor network, potentially including central pattern generating circuits (CPGs). In response, motoneurons decrease their excitable properties, thus maintaining normal locomotor output to the body wall muscles. Nevertheless, changes in the premotor network lead to a reduction in activity wave progression and thus reduced larval crawling speed, also manifesting a reduction in overall network stability.

### NMJ structural development is affected by CP manipulations, but not synaptic transmission

Despite embryonic heat stress leading to significant NMJ structural changes, in terms of presynaptic terminal overgrowth (an average increase of active zone number by 30%) and overall reduction of GluRIIA (by 50%), evoked transmission resulting from the firing of single action potentials in the motor axon remained normal. Distal boutons had previously been shown to house release sites that are preferentially recruited under low frequency stimulation regimes, such as used for making NMJ recordings (Peled and Isacoff, 2011). Speculatively, preferential maintenance of GluRIIA in distal boutons, might have explained maintenance of normal synaptic transmission at low frequency stimulation regimes. However, we found no evidence for this. On the contrary, in embryonic heat stress-manipulated specimens, GluRIIA levels were comparably reduced in both proximal and distal boutons (data not shown). An alternative explanation for maintained synaptic transmission in the face of increased active zone number is a reduction in release probability. Although we did not conduct functional imaging to investigate this directly, an indirect indication for this is the increased facilitation in animals that had experienced embryonic heat stress, which indicates a reduction in presynaptic release probability across the whole NMJ (Fig. 4D).

On reflection, to find evoked synaptic transmission at the NMJ not significantly affected by CP manipulations, is perhaps not unexpected. A well-established feature of this particular synapse is that its transmission is tightly regulated so as to maintain constancy through the deployment of multiple NMJ plasticity mechanisms, including presynaptic homeostatic potentiation (PHP), which we have shown to function effectively even after an embryonic heat stress manipulation (Fig. 4G) (Delvendahl and Müller, 2019; Frank et al., 2020, 2006; Goel and Dickman, 2021).

### Distinct developmental windows with sensitivity to heat stress suggest sequential network maturation

It is unlikely that structural changes at the NMJ contribute to making the nervous system less stable, or that they effect CP-regulated changes to larval behaviour, such as crawling speed or social feeding (Williamson et al., 2021). Based on previous CP manipulations targeted to various neuronal subsets (Giachello et al., 2021), it seemed more likely that behavioural phenotypes, including reduced crawling speed and increased recovery times from electroshock-induced seizures, would be the consequence of changes in the central circuitry. Indeed, we found that the developing body wall muscles have an earlier CP that is sensitive to heat stress (13-16 hrs AEL) and leads to changes in postsynaptic GluR composition (Fig. 3), than the central locomotor circuitry (17-19 hrs AEL) (Figs, 1 & 8A). By applying heat stress manipulations to these distinct developmental phases we were able to demonstrate that behavioural phenotypes on both larval crawling speed and network stability are linked to CP-induced changes to the central circuitry, not the peripheral NMJ (Figs. 1, 3 & 8A). These CPs correlate with the developmental phases when the body wall muscles and central neurons become electrically active, respectively (Broadie and Bate, 1993; Baines and Bate, 1998). Thus, they reflect the sequence of earlier myogenic and subsequent neuronal maturation (Crisp et al., 2008). This is comparable with observations from mammalian systems, where different parts of the developing brain mature in sequence (Hensch, 2005; Hensch and Quinlan, 2018; Reh et al., 2020).

Mechanistically, heat stress causes increased neuronal activity (Oswald et al., 2018; Sigrist et al., 2003; Zhong and Wu, 2004). Functional imaging of motoneurons showed that in larvae acute temperature elevations cause corresponding increases in both fictive locomotor activity waves (Fig. 1B) as well as larval crawling (Fig. 8B). Interestingly, this manipulation, when carried out during the embryonic CP, leads to a locomotor network with significantly slower intersegmental propagation of activity waves and, similarly, reduced crawling speed (Fig. 7). One possibility is that heat stress during the CP might cause a lasting malfunctioning of pacemaker activity. Alternatively, the slower central pattern of locomotor network activity in larval stages could be the consequence of a network-wide homeostatic response to an abnormally fast rate provoked by transient heat stress during the embryonic CP. The latter explanation is compatible with our recent findings of the CP as an integration window of excitatory *vs* inhibitory network activity (Hunter et al., 2024) and the identification of nitric oxide signalling pathways downstream of CP network activity (Giachello et al., 2021). To identify additional genetic pathways associated with CP plasticity, we propose that the NMJ, as a large, readily accessible and morphometrically quantifiable peripheral synapse, could serve as a convenient and effective assay.

## Acknowledgements

This work was made possible through support by a Walter Benjamin Programme Fellowship to NK by the Deutsche Forschungsgemeinschaft (DFG) KR5597/1-1, and funding from the Biotechnology and Biological Sciences Research Council (BBSRC) to ML (BB/V014943/1) and the Wellcome Trust, through a Joint Wellcome Trust Investigator Award to RAB and ML (217099/Z/19/Z). Research reported in this publication was supported by an institutional startup fund from Texas A&M University (AAZ). DSC was supported by the European Molecular Biology Organization (EMBO) with a long-term EMBO fellowship (ALTF 62-2021). The work benefited from the Imaging Facility, Department of Zoology, supported by Matt Wayland and Tom Pettini, and funds from a Wellcome Trust Equipment Grant (WT079204) with contributions by the Sir Isaac Newton Trust in Cambridge, including Research Grant [18.07ii(c)]. Work on this project further benefited from the Manchester Fly Facility, established through funds from the University and the Wellcome Trust (grant 087742/Z/08/Z).

The authors would like to thank members of the Baines and Landgraf research teams for feedback on the manuscript. The authors are grateful to Ela Serpe for Rabbit polyclonal anti-GluRIIB antiserum.

Stocks obtained from the Bloomington Drosophila Stock Center (NIH P40OD018537) were used in this study.

The 8B4D2 (MH2B) monoclonal antibody to visualise GluRIIA, developed C. Goodman, was obtained from the Developmental Studies Hybridoma Bank, created by the NICHD of the NIH and maintained at The University of Iowa, Department of Biology, Iowa City, IA 52242.

## Materials and Methods

### Fly husbandry

Wild-type and transgenic strains were maintained on standard yeast–agar–cornmeal medium at 25°C. For CP manipulations, animals were kept in laying pots with apple juice agar plates and yeast paste for 6h. Collected eggs were either kept at 25°C for controls or at 32°C for 24h, before being moved back to 25°C until animals developed to the L3 larval stage.

### Dissection and Immunostainings

Wandering L3 larvae were dissected in Sørensen’s sodium phosphate buffered saline (0.1 M, pH 7.2) and fixed for 10min at room temperature with Bouińs fixative (Sigma-Aldrich). After fixing, specimens were washed in washing solution (Sørensen’s buffer + 0.25% BSA + 0.3% TritonX-100 from Sigma-Aldrich) and incubated with mouse nc82 (anti-Bruchpilot to reveal active zones) antibody (1:100, DSHB) overnight, visualized with goat α-mouse StarRed (1:1000, Abberior, cat.no.STRED-1001-500UG, Lot. 2042PK-1) and AlexaFluor488 conjugated goat α-HRP (1:1000, Jackson ImmunoResearch, cat.no. 123-545-021, Lot. 152767) to visualise presynaptic, neuronal structures. For stainings of the postsynaptic glutamate receptor subunits GluRIIA and GluRIIB we used monoclonal mouse-α-DGluRIIA antibody (1:100, DSHB, product ID: 8B4D2 (MH2B)) and polyclonal rabbit-α-DGluRIIB antiserum (1:1000, gift from Mihaela Serpe (Sulkowski et al., 2016)) in conjunction with goat α-mouse StarRed (1:1000) and goat α-rabbit Atto594 secondary antibodies (1:1000, Hypermol, cat.no. 2306, Lot. 0356RG22), respectively, in combinations with AlexaFluor488 conjugated goat α-HRP (1:1000, Jackson ImmunoResearch, cat.no. 123-545-021, Lot. 152767) to visualise presynaptic terminals.

### Fly stocks and other Key Resources

| Key Resources Table |  |  |  |  |
| --- | --- | --- | --- | --- |
| Reagent type (species) or resource | Designation | Source or reference | Identifiers | Additional information |
| strain, strain background (*Drosophila melanogaster*) | Oregon-R | Bloomington Drosophila Stock Center | RRID:BDSC_2376<br>FLYB: FBst0002376 | wildtype control |
| strain, strain background (*Drosophila melanogaster*) | Canton S | Bloomington Drosophila Stock Center | RRID:BDSC_64349<br>FLYB: FBst0064349 | wildtype control |
| genetic reagent (*D. melanogaster*) | GMR94G06-Gal4 | Bloomington Drosophila Stock Center | RRID:BDSC_40701<br>FLYB: FBti0139793 | FlyBase symbol: w1118; ; GMR94G06-GAL4 [attP2] |
| genetic reagent (*D. melanogaster*) | A27h-Gal4 | Bloomington Drosophila Stock Center | RRID:BDSC_49939<br>FLYB: FBti0135233 | FlyBase symbol: w1118; ; P{y[+t7.7]w[+mC]=GM R36G02-GAL4}attP2 |
| genetic reagent (*D. melanogaster*) | A27h-split-Gal4 | Created by A.A. Zarin, this study. Components available in Bloomington Drosophila Stock Center | RRID:BDSC_70953<br>FLYB: FBti0188221<br>RRID:BDSC_68768<br>FLYB: FBti0191234 | FlyBase symbol: w1118; 34F03-AD [attP40] ; 36G02-DBD [attP2] |
| genetic reagent (*D. melanogaster*) | 20XUAS-IVS-jGCaMP8f | Bloomington Drosophila Stock Center | RRID:BDSC_92587<br>FLYB: FBti0215881 | FlyBase symbol: w1118; ; 20XUAS-IVS-jGCaMP8f [VK00005] |
| genetic reagent<br>(*D. melanogaster*) | 20XUAS-Chronos-mVenus | Bloomington<br>Drosophila<br>Stock Center | RRID:BDSC_77115<br>FLYB:FBti0195682 | FlyBase<br>symbol:<br>w1118;<br>20XUAS-Chronos-mVenus<br>[attP40] |
| antibody | anti-Bruchpilot<br>(NC82)<br>(Mouse<br>monoclonal) | DSHB | Cat# nc82,<br>RRID:AB_2314866 | IF(1:100) |
| antibody | anti-DGluRIIA<br>8B4D2<br>(MH2B)<br>(Mouse<br>monoclonal) | DSHB | Cat# 8B4D2<br>(MH2B),<br>RRID:<br>AB_528269 | IF(1:100) |
| antibody | anti-DGluRIIB<br>(Rabbit<br>polyclonal) | Mihaela<br>Serpe<br>(Sulkowski et<br>al., 2016) |  | IF(1:1000) |
| antibody | anti-mouse-STAR Red<br>(Goat<br>polyclonal) | Abberior | Cat#:<br>STRED-1001-500UG;<br>RRID:AB_3068620 | IF(1:1000) |
| antibody | anti-rabbit-ATTO594<br>(Goat<br>polyclonal) | Hypermol | Cat#: 2306; | IF(1:1000) |
| antibody | Alexa<br>Fluor488<br>Goat Anti-Horseradish<br>Peroxidase<br>(Goat<br>polyclonal) | Jackson<br>ImmunoRese<br>arch | Cat#: 123-545-021;<br>AB_2338965 | IF(1:1000) |

### Image acquisition

Confocal images were acquired, using a Leica Stellaris point scanning confocal microscope, with an 40x/1.25N.A. oil immersion lens (Leica) and a 1.25 digital zoom. Images were acquired with a voxel size 0.16x 0.16x 0.3 µm. Images were processed using FIJI and Affinity Designer.

### Two electrode voltage clamp

Two Electrode Voltage Clamp (TEVC) recordings were performed in HL3.1 saline (Feng et al., 2004) composition in mM: NaCl 70, KCl 2.5, MGCl_2_ 4, CaCl_2_ 2 (if not other mentioned), NaHCO_3_ 10, trehalose 5, sucrose 115, HEPES 5 (pH was adjusted to 7.24-7.25 with 1M NaOH), using an Olympus BX50WI compound microscope with 60X dipping (Olympus 60x/0.9 N.A.) objective lenses. Experiments were performed on third-instar larval NMJs of the dorsal acute muscle 1 (DA1) in segments A3 and A4. The respective nerve was sucked in with a self-built suction electrode (Johnson et al., 2007) which was filled with HL3.1 saline. Subsequently, the nerve was stimulated for 0.1ms to trigger action potential firing. Briefly, voltage was stepwise increased until the first synaptic transmission could be recorded. From that voltage, 1 V was added to reliably evoke APs in control conditions, in 32°C CP manipulated animals 2-3 V had to be added to reliably recruit aCC and RP2. For PhTx experiments, samples were incubated in 4 µM PhTx final concentration (Frank et al., 2006) for 10 min. While dissecting, great care was taken to not stretch animals too much. Successful PhTx application was determined by dividing miniature EPSC amplitude of PhTx treated animals by the average Amplitude of control mEPSCs. If PhTx treated mEPSCs were at least 40% reduced compared to control, PhTx application was considered successful (Frank et al., 2006). Quantal content was determined by dividing EPSC by mEPSC for each recording. Recordings were only further considered from cells with an initial Vm below -50 mV and input resistance between 8-10 MΩ, using intracellular electrodes filled with 3M KCl with resistances of ∼20 MΩ (Electrodes for current injection and the membrane voltage). Current passing und recording electrodes were pulled from Borosilicate glass microelectrodes with filament (OD1 mm, ID 0.58mm; Harvard Apparatus LTD, GC100F-10 Part No. 30-0019), suction electrodes from Borosilicate glass microelectrodes without filament (OD1 mm, ID 0.58mm; Harvard Apparatus LTD, GC100-10 Part No. 30-0016) with a Flaming/Brown Puller Model P-97 (Sutter Instruments USA). Excitatory postsynaptic currents (EPSCs) were evoked by an Isolated Pulse Stimulator (Model 2100 A-M Systems USA), recorded at a clamped voltage of -60 mV with a sampling rate of 50 kHz and filtered with the anti-alias filter at 25 kHz using the AxoClamp 2B amplifier (Molecular Devices Axon Instruments USA). Data analysis was done with Clampfit 10.7. Traces were first filtered with a 360 Hz Gaussian low pass filter, then template searches were run over the trace for single pulses and spontaneous miniature excitatory postsynaptic currents (mEPSCs or minis). The resulting files were used to analyse mEPSC and EPSC properties. Rise and decay tau were measured with an one-term exponential product fit (Levenberg-Marquadt method with a max of 5000 iterations)

### Current clamp muscle recordings

Current clamp recordings from muscle DA1 were performed in HL3.1 saline (see above), in semi intact, fillet dissected wandering L3 larvae with intact nervous system. After finishing the dissection, only animals performing crawling-like movements were considered for further experiments. To block muscle contractions, a 500 µM stock solution of nifedipine in HL3.1 and 1% DMSO was added to the bath for a final concentration of 50 µM nifedipine and 0.1% DMSO for 15 min. Recordings were only further considered from cells with an initial Vm below -60 mV and input resistance between 8-10 MΩ, using intracellular electrodes filled with 3M KCl with resistances of ∼20 MΩ. Spontaneous activity was recorded for 5min. Burst analysis was performed using the Clampfit Burst Analysis, using the Poisson Surprise model with a minimum of 5 events per burst.

### Electroshock assay

A handheld stimulator consisting of two tungsten wires (0.1 mm diameter) fixed to a plastic rod was used. Wires were shaped to be ∼1-2 mm apart, attached to a DS2A Mk.II stimulator (Digitimer Ltd), which was set to provide a 6 V, 2 s current pulse. Larvae were placed on a plastic plate lid, with tissue paper used to dry residual moisture from the larvae. Once larvae had returned to normal crawling, the tungsten wires were brought into contact with the larvae, positioned on the larval cuticle, perpendicular to and over the approximate position of the CNS, with gentle pressure. Current stimulation was then delivered, which usually induced a sustained contraction and paralysis representing a seizure (videos can be viewed in Huertas Radi et al., 2025 and Marley and Baines 2011). The time to resumption of normal behaviour (regular, whole body forward propagating peristaltic waves) was recorded and defined as the duration of recovery from seizure.

### Dissection and CNS mounting for calcium imaging and electrophysiology

Larval CNSs were dissected and mounted for electrophysiological recordings and for GCaMP imaging. Third instar (L3) larvae (of either sex) were dissected in a dish in standard saline (135 mM NaCl (Fisher Scientific), 5 mM KCl (Fisher Scientific), 4 mM MgCl_2_·6H_2_O (Sigma-Aldrich), 2mM CaCl_2_·2H_2_O (Fisher Scientific), 5 mM TES (Sigma-Aldrich), 36 mM sucrose (Fisher Scientific), pH 7.15), to remove the CNS (ventral nerve cord and brain lobes). The isolated CNS was then transferred to a droplet of external saline, within which it was laid flat (dorsal side up) and glued (GLUture Topical Tissue Adhesive; World Precision Instruments USA) to a Sylgard-coated cover slip (1 to 2mm depth of cured SYLGARD Elastomer (Dow-Corning USA) on a 22 x 22mm square coverslip). The preparation was then placed on a glass slide and viewed under a microscope (Olympus BX51-WI).

### Calcium imaging

Dissected CNSs were imaged (QImaging EXi-Aqua; photometrics, Arizona, US) with an acquisition frame rate of 10Hz and frame duration of 100 ms using a x20 immersion lens. GCaMP8f was excited using a 470 nm collimated LED (Thorlabs, New Jersey, US). Time-course of neuronal fluorescence was determined in WinFluor (V.4.1.5; University of Strathclyde, UK) by adding regions of interest (ROIs) to axons, close to the cell body (Streit et al, 2016). In Clampfit (10.3.1.5; Molecular Devices, California, US) fluorescence traces were smoothed using a lowpass BoxCar filter using 11 smoothing points prior to analysis.

For motoneuron calcium transients (Fig. 1), recordings were conducted for 600s, whilst temperature of the explanted CNS was increased steadily from ∼23°C to ∼34°C using a digital heating unit (CO 102; Linkam Scientific, Surrey, UK). For sham recordings, temperature was maintained at a constant 25°C. Temperature was measured in Clampex (10.3.1.5; Molecular Devices, California, US) using a temperature sensor placed in the external saline surrounding the brain, via a temperature controller (TC-10; npi electronic GmbH, Tamm, Germany). The frequency of calcium fluorescence peaks was measured for each segment (A8/9 – A1) in three sequential 200 s time periods and averaged per larva. Average temperature was calculated for each recording during these same 200 s time periods.

For calcium imaging of interneurons after CP manipulation (Fig. 5), 180 s recordings were taken. Time lag was determined by manually assessing the time between peak fluorescence corresponding to ipsilaterally adjacent interneurons (segments A6 – A1. For each recording the first complete forward waves of activity were used (up to three repeats per trace) to generate an average intersegment duration for each pair of interneurons.

### Whole cell patch clamp electrophysiology

CNSs were dissected, mounted (see calcium imaging methods) and viewed under a 60x water-immersion lens. To access soma, 1% protease (Streptomyces griseus, Type XIV, Sigma-Aldrich, in external saline) contained within a wide-bore glass pipette (GC100TF-10; Harvard Apparatus UK, approx. 10 µm opening) was applied to abdominal segments, roughly between A5-A2 (Baines and Bate, 1998). This was done to remove overlaying glia to facilitate access to underlying nerve cell soma (e.g., aCC). Motoneurons were identified by anatomical position and relative cell size, with aCC being positioned close to the midline and containing both an ipsilateral and contralateral projection. Whole-cell patch recordings were made using borosilicate glass pipettes (GC100F-10, Harvard Apparatus) that were fire polished to resistances of 10 - 15MΩ when filled with intracellular saline (140 mM potassium-D-gluconate (Sigma-Aldrich), 2 mM MgCl_2_·6H_2_O (Sigma-Aldrich), 2 mM EGTA (Sigma-Aldrich), 5 mM KCl (Fisher Scientific), and 20 mM HEPES (Sigma-Aldrich), (pH 7.4). Input resistance was measured in the ‘whole cell’ configuration, and only cells that had an input resistance ≥ 0.5 GΩ were used for experiments. Cell capacitance and break-in resting membrane potential was also measured for each cell recorded. Data for current steps recordings was captured using a Multiclamp 700B amplifier controlled by pCLAMP (version 10.7.0.3), via an analogue-to-digital converter (Digidata 1440A, Molecular Devices). Trace/s were sampled at 20 kHz and filtered online at 10 kHz.

### Current step recordings

Mecamylamine (200 µM in saline) was used to block postsynaptic nACh receptors to synaptically isolate neurons for experiments designed to measure intrinsic motoneuron excitability. Once patched, neurons were brought to a membrane potential of -60 mV using current injection. Each recording consisted of 20 x 4 pA (500 ms) current steps, including an initial negative step, giving a range of -4 to +72 pA. Number of spikes were counted and plotted against injected current. Both cell capacitance and input resistance were compared between conditions to ensure that any observed differences in excitability were not due to differences in either cell size or resistance.

### Synaptic drive to motoneurons

To measure synaptic drive between A27h premotor interneurons and aCC, A27h neurons were identified via *Chronos-mVenus* reporter (as above), allowing corresponding aCC motoneurons to be patched. Recordings were conducted in current-clamp, with current injected into aCC sufficient to evoke a mean spike frequency of ∼10-20 Hz, prior to optogenetic stimulation. Recordings consisted of: individual sweeps, beginning with 1 s LED off, followed by 1 s LED on, and 1 s LED off (repeated 5-8 times per cell). Change in AP frequency (%) was calculated for each recording (per sweep) comparing AP frequency before and during stimulation, which was then averaged for each cell.

Spontaneous rhythmic currents (SRCs) were recorded for 180 s with aCC voltage clamped at -60 mV. Amplitude, duration, and frequency were measured for the first 15 events of each trace using Clampfit’s event detection, threshold search, function. The first 15 events with a stable baseline, an amplitude of >300 pA and no double peaks were accepted for analysis. Amplitude was measured as the change from baseline immediately prior to event initiation to the peak current and normalised for cell capacitance, providing a measure of current density. SRC duration was measured as the width of an event at 100 pA amplitude.

### Spontaneous motoneuron recordings

Cell-attached recordings, measuring motoneuron activity, were conducted using unpolished patch electrodes. Motoneuron spontaneous activity was recorded using an adapted cell-attached current clamp protocol (Multiclamp 700B amplifier with I=0) to measure passive action potential bursting properties. In Clampfit (version 10.3.1.5), 180 s recordings, were analysed to detect action potential ‘spike times’. These were imported into MATLAB and analysed using a custom script (developed by Mituzaite et al., 2021). An action potential burst was considered as an event consisting of at least 3 spikes occurring within 100 ms and ending when no spike was detected for 100 ms.

### Crawling Assay

Crawling behaviour was recorded at the mid-third instar stage, 72 hours after larval hatching (ALH). Larvae were rinsed in water and placed inside a 24cm x 24cm crawling arena with a base of 5mm thick 0.8% agarose gel (Bacto Agar), situated inside an incubator. Temperature was maintained at 25±0.5°C, reported via a temperature probe in the agar medium. Humidity was kept constant. Larval crawling was recorded using a frustrated total internal reflection-based imaging method (FIM) in conjunction with the tracking software FIMTrack (Risse et al., 2017, 2013) using a Basler acA2040-180km CMOS camera, fitted with a 16mm KOWA-IJM3sHC.SW-VIS-NIR lens, acquisition controlled by Pylon software (Basler) and Streampix (v.6) software (NorPix). Larvae were recorded for 20 minutes at five frames per second.

Recordings were split into four 5-minute sections with the first five minutes of acclimatisation discarded. The remaining three 5-minute sections were used to analyse crawling speed, choosing crawling periods of uninterrupted forward crawling, devoid of pauses, turning or collision events. Each larva was sampled up to once per 5-minute section, i.e. up to three data points were derived per larva from up to three consecutive 5-minute segments. Crawling speed was calculated using FIMTrack software.

### Statistics

Statistical analyses were done using GraphPad Prism Software (Version 10.1.2). Datasets were tested for normal distribution with the Shapiro-Wilk Test. Normally distributed data were then tested with students t-test (for pairwise comparison). Normally distributed analysis for more than two groups was done with a one-way ANOVA and post hoc tested with a Tukey multiple comparison test. Non-normally distributed data sets of two groups were tested with Mann-Whitney U Test (pairwise comparison) and datasets with more than two groups were tested with a Kruskal Wallis ANOVA and post hoc tested with a Dunns multiple post hoc comparison test. For whole cell electrophysiology experiments in isolated neurons, linear regression analysis was used to compare the intercepts between conditions. This analysis was restricted to linear sections of data, represented by inset graphs. For all datasets mean and standard error of mean (SEM) are shown. Significance levels were * p<0.05; ** p<0.01; *** p<0.001; **** p<0.0001. Sample sizes are stated in figures or legends, each data point derived from a unique specimen, unless otherwise stated; e.g. Figure 3B, NMJ sizes were sampled at up to four data points per larva; for larval crawling speeds up to three data points were derived per larva, one for each unique 5-minute segment of recorded crawling. All experiments indiscriminately included larvae of both sexes.

## References

Ackerman SD, Perez-Catalan NA, Freeman MR, Doe CQ. Astrocytes close a motor circuit critical period. 2021. Nature 592:414–420. doi:10.1038/s41586-021-03441-2

Ashburner M, Carson HL, Thompson JN. 1983. Ectophysilogy: Abiotic FactorsThe Genetics and Biology of Drosophila,. pp. 105–170.

Baines RA, Bate M. 1998. Electrophysiological development of central neurons in the Drosophila embryo. J Neurosci. 18:4673–83. doi: 10.1523/JNEUROSCI.18-12-04673.1998.

Baines RA, Robinson SG, Fujioka M, Jaynes JB, Bate M. 1999. Postsynaptic expression of tetanus toxin light chain blocks synaptogenesis in Drosophila. Current Biology 9:1267–S1. doi:10.1016/S0960-9822(99)80510-7

Baines RA, Uhler JP, Thompson A, Sweeney ST, Bate M. 2001. Altered Electrical Properties in Drosophila Neurons Developing without Synaptic Transmission. The Journal of Neuroscience 21:1523–1531. doi:10.1523/JNEUROSCI.21-05-01523.2001

Bate M, Rushton E, Frasch M. A dual requirement for neurogenic genes in Drosophila myogenesis. Dev Suppl. 1993:149–61. PMID: 8049469.

Baumann NS, Sears JC, Broadie K. 2024. Experience-dependent MAPK/ERK signaling in glia regulates critical period remodeling of synaptic glomeruli. Cell Signal 120:111224. doi:10.1016/j.cellsig.2024.111224

Broadie KS, Bate M. 1993. Development of larval muscle properties in the embryonic myotubes of Drosophila melanogaster. J Neurosci. 13:167–80. doi: 10.1523/JNEUROSCI.13-01-00167.1993

Carreira-Rosario A, York RA, Choi M, Doe CQ, Clandinin TR. 2021. Mechanosensory input during circuit formation shapes Drosophila motor behavior through patterned spontaneous network activity. Current Biology 31:5341–5349.e4. doi:10.1016/j.cub.2021.08.022

Chodankar A, Sadanandappa MK, VijayRaghavan K, Ramaswami M. 2020. Glomerulus-Selective Regulation of a Critical Period for Interneuron Plasticity in the *Drosophila* Antennal Lobe. The Journal of Neuroscience 40:5549–5560. doi:10.1523/JNEUROSCI.2192-19.2020

Corke J, Huertas Radi M, Landgraf M, Baines RA. 2025. Maturation of GABAergic signalling times the opening of a critical period in Drosophila melanogaster. doi:10.1101/2025.04.24.650400

Coulson B, Hunter I, Doran S, Parkin J, Landgraf M, Baines RA. 2022. Critical periods in Drosophila neural network development: Importance to network tuning and therapeutic potential. Front Physiol 13:1–11. doi:10.3389/fphys.2022.1073307

Crisp S, Evers JF, Fiala A, Bate M. 2008. The development of motor coordination in Drosophila embryos. Development 135:3707–3717. doi:10.1242/dev.026773

Crisp SJ, Evers JF, Bate M. 2011. Endogenous Patterns of Activity Are Required for the Maturation of a Motor Network. Journal of Neuroscience 31:10445–10450. doi:10.1523/JNEUROSCI.0346-11.2011

Delvendahl I, Müller M. 2019. Homeostatic plasticity—a presynaptic perspective. Curr Opin Neurobiol 54:155–162. doi:10.1016/j.conb.2018.10.003

Dombrovski M, Condron B. 2021. Critical periods shaping the social brain: A perspective from Drosophila. BioEssays 43:2000246. doi:10.1002/bies.202000246

Dyson A, Ryan M, Garg S, Evans DG, Baines RA. 2022. Loss of NF1 in Drosophila Larvae Causes Tactile Hypersensitivity and Impaired Synaptic Transmission at the Neuromuscular Junction. Journal of Neuroscience 42:9450–9472. doi:10.1523/JNEUROSCI.0562-22.2022

Feng Y, Ueda A, Wu CF. 2004. A modified minimal hemolymph-like solution, HL3.1, for physiological recordings at the neuromuscular junctions of normal and mutant Drosophila larvae. J Neurogenet 18:377–402. doi:10.1080/01677060490894522

Frank CA, James TD, Müller M. 2020. Homeostatic control of Drosophila neuromuscular junction function. Synapse 74:22133. doi:10.1002/syn.22133

Frank CA, Kennedy MJ, Goold CP, Marek KW, Davis GW. 2006. Mechanisms Underlying the Rapid Induction and Sustained Expression of Synaptic Homeostasis. Neuron 52:663–677. doi:10.1016/j.neuron.2006.09.029

Fujioka M, Lear BC, Landgraf M, Yusibova GL, Zhou J, Riley KM, Patel NH, Jaynes JB. 2003. Even-skipped, acting as a repressor, regulates axonal projections in Drosophila. Development 130:5385–5400. doi:10.1242/dev.00770

Fushiki A, Kohsaka H, Nose A. 2013. Role of Sensory Experience in Functional Development of Drosophila Motor Circuits. PLoS One 8:e62199. doi:10.1371/journal.pone.0062199

Genç Ö, Davis GW. 2019. Target-wide Induction and Synapse Type-Specific Robustness of Presynaptic Homeostasis. Current Biology 29:3863–3873.e2. doi:10.1016/j.cub.2019.09.036

Giachello CNG, Baines RA. 2015. Inappropriate Neural Activity during a Sensitive Period in Embryogenesis Results in Persistent Seizure-like Behavior. Current Biology 25:2964–2968. doi:10.1016/j.cub.2015.09.040

Giachello CNG, Fan YN, Landgraf M, Baines RA. 2021. Nitric oxide mediates activity-dependent change to synaptic excitation during a critical period in Drosophila. Sci Rep 11:1–12. doi:10.1038/s41598-021-99868-8

Giachello CNG, Fan YN, Landgraf M, Baines RA. 2019. Activity manipulation of an excitatory interneuron, during an embryonic critical period, alters network tuning of the Drosophila larval locomotor circuit. bioRxiv. doi:10.1101/780221

Giachello CNG, Zarin AA, Kohsaka H, Fan YN, Nose A, Landgraf M, Baines RA. 2020. Electrophysiological validation of premotor interneurons monosynaptically connected to the aCC motoneuron in the Drosophila larval CNS. bioRxiv 1–19. doi:10.1101/2020.06.17.156430

Goel P, Dickman D. 2021. Synaptic homeostats: latent plasticity revealed at the Drosophila neuromuscular junction. Cellular and Molecular Life Sciences. doi:10.1007/s00018-020-03732-3

Hageter J, Starkey J, Horstick EJ, Hageter J, Starkey J, Horstick EJ. 2023. Article Thalamic regulation of a visual critical period and motor behavior ll ll Thalamic regulation of a visual critical period and motor behavior. CellReports 42:112287. doi:10.1016/j.celrep.2023.112287

Harrison R V, Gordon KA, Mount RJ. 2005. Is There a Critical Period for Cochlear Implantation in Congenitally Deaf ChildrenAnalyses of Hearing and Speech Perception Performance after Implantation. Dev Psychobiol 46:155–292. doi:DOI 10.1002/dev.20052

Heckman EL, Doe CQ. 2022. Presynaptic contact and activity opposingly regulate postsynaptic dendrite outgrowth. Elife 11. doi:10.7554/eLife.82093

Hensch TK. 2005. Critical period plasticity in local cortical circuits. Nat Rev Neurosci. doi:10.1038/nrn1787

Hensch TK, Quinlan EM. 2018. Critical periods in amblyopia. Vis Neurosci. doi:10.1017/S0952523817000219

Hess EH. 1975. Imprinting: Early experience and the dvelopmental psychobiology of attachment. Psychosom Med 37:188–190.

Horn G. 2004. Pathways of the past: the imprint of memory. Nat Rev Neurosci 5:108–121. doi:10.1038/nrn1324

Hubel DH, Wiesel TN. 1970. The period of susceptibility to the physiological effect of unilateral eye closure in kittens. Journal of Physiology 206:419–436. doi:10.1113/jphysiol.1970.sp009022

Huertas Radi M, Corke J, Baines RA. Seizure Activity Induced by Electroshock in Drosophila Larvae. J Vis Exp. 2025 Jun 6;(220). doi: 10.3791/68431. PMID: 40549583.

Hunter I, Coulson B, Pettini T, Davies JJ, Parkin J, Landgraf M, Baines RA. 2024. Balance of activity during a critical period tunes a developing network. Elife 12:1–23. doi:10.7554/eLife.91599

Hunter I, Coulson B, Zarin AA, Baines RA. 2021. The Drosophila Larval Locomotor Circuit Provides a Model to Understand Neural Circuit Development and Function. Front Neural Circuits. doi:10.3389/fncir.2021.684969

Johnson BR, Hauptman SA, Bonow RH. 2007. Construction of a Simple Suction Electrode for Extracellular Recording and Stimulation. Journal of Undergraduate Neuroscience Education 6:A21.

Jusyte M, Blaum N, Böhme MA, Berns MMM, Bonard AE, Vámosi ÁB, Pushpalatha K V., Kobbersmed JRL, Walter AM. 2023. Unc13A dynamically stabilizes vesicle priming at synaptic release sites for short-term facilitation and homeostatic potentiation. Cell Rep 42. doi:10.1016/j.celrep.2023.112541

Kiral FR, Dutta SB, Linneweber GA, Hilgert S, Poppa C, Duch C, von Kleist M, Hassan BA, Hiesinger PR. 2021. Brain connectivity inversely scales with developmental temperature in Drosophila. Cell Rep 37:110145. doi:10.1016/j.celrep.2021.110145

Kratschmer P, Lowe SA, Buhl E, Chen K, Kullmann DM, Pittman A, Hodge JJL, Jepson JEC. 2021. Impaired Pre-Motor Circuit Activity and Movement in a Drosophila Model of KCNMA1 -Linked Dyskinesia. Movement Disorders 36:1158–1169. doi:10.1002/mds.28479

Kuntz SG, Eisen MB. 2014. Drosophila embryogenesis scales uniformly across temperature in developmentally diverse species. PLoS Genet. 10(4):e1004293. doi: 10.1371/journal.pgen.1004293.

Landgraf M, Jeffrey V, Fujioka M, Jaynes JB, Bate M. 2003. Embryonic Origins of a Motor System: Motor Dendrites Form a Myotopic Map in Drosophila. PLoS Biol 1:e41. doi:10.1371/journal.pbio.0000041

Lee Y, Montell C. 2013. Drosophila TRPA1 functions in temperature control of circadian rhythm in pacemaker neurons. Journal of Neuroscience 33:6716–6725. doi:10.1523/JNEUROSCI.4237-12.2013

Leier HC, Foden AJ, Jindal DA, Wilkov AJ, Van der Linden Costello P, Vanderzalm PJ, Coutinho-Budd J, Tabuchi M, Broihier HT. 2025. Glia control experience-dependent plasticity in an olfactory critical period. Elife 13. doi:10.7554/eLife.100989

Lowe SA, Wilson AD, Aughey GN, Banjaree A, Goble T, Simon-Batsford N, Sanderson A, Kratschmer P, Balogun M, Gao H, Aw SS, Jepson JEC. 2023. BK channel gain-of-function disrupts limp control by suppressing neurotransmission during a critical developmental window. bioRxiv. doi:10.1101/2023.09.20.558625

Marder E. 2023. Individual Variability, Statistics, and the Resilience of Nervous Systems of Crabs and Humans to Temperature and Other Perturbations. eNeuro 10:ENEURO.0425-23.2023. doi:10.1523/ENEURO.0425-23.2023

Marder E, Rue MCP. 2021. From the Neuroscience of Individual Variability to Climate Change. The Journal of Neuroscience 41:10213–10221. doi:10.1523/JNEUROSCI.1261-21.2021

Mòdol L, Moissidis M, Selten M, Oozeer F, Marín O. 2024. Somatostatin interneurons control the timing of developmental desynchronization in cortical networks. Neuron 112:2015–2030.e5. doi: 10.1016/j.neuron.2024.03.014

Müller M, Davis GW. 2012. Transsynaptic control of presynaptic Ca 2+ influx achieves homeostatic potentiation of neurotransmitter release. Current Biology 22:1102–1108. doi:10.1016/j.cub.2012.04.018

Nardou R, Sawyer E, Song YJ, Wilkinson M, Padovan-Hernandez Y, de Deus JL, Wright N, Lama C, Faltin S, Goff LA, Stein-O’Brien GL, Dölen G. 2023. Psychedelics reopen the social reward learning critical period. Nature 618:790–798. doi:10.1038/s41586-023-06204-3

Nelson N, Vita DJ, Broadie K. 2024. Experience-dependent glial pruning of synaptic glomeruli during the critical period. Sci Rep 14:9110. doi:10.1038/s41598-024-59942-3

Oswald MCW, Brooks PS, Zwart MF, Mukherjee A, West RJH, Giachello CNG, Morarach K, Baines RA, Sweeney ST, Landgraf M. 2018. Reactive oxygen species regulate activity-dependent neuronal plasticity in Drosophila. Elife 7. doi:10.7554/eLife.39393

Peled ES, Isacoff EY. 2011. Optical quantal analysis of synaptic transmission in wild-type and rab3-mutant Drosophila motor axons. Nat Neurosci 14:519–526. doi:10.1038/nn.2767

Pérez-Moreno JJ, O’Kane CJ. 2019. GAL4 Drivers Specific for Type Ib and Type Is Motor Neurons in Drosophila. G3 Genes|Genomes|Genetics 9:453–462. doi:10.1534/g3.118.200809

Powsner L. 1935. The Effects of Temperature on the Durations of the Developmental Stages of Drosophila melanogaster. Physiol Zool 8:474–520. doi:10.1086/physzool.8.4.30151263

Pulver SR, Bayley TG, Taylor AL, Berni J, Bate M, Hedwig B. Imaging fictive locomotor patterns in larval Drosophila. 2015. J Neurophysiol 114:2564–77. doi: 10.1152/jn.00731.2015

Reh RK, Dias BG, Nelson CA 3rd, Kaufer D, Werker JF, Kolb B, Levine JD, Hensch TK. 2020. Critical period regulation across multiple timescales. Proc Natl Acad Sci U S A 117:23242–23251. doi: 10.1073/pnas.1820836117

Risse B, Berh D, Otto N, Klämbt C, Jiang X. 2017. FIMTrack: An open source tracking and locomotion analysis software for small animals. PLoS Comput Biol 13:1–15. doi:10.1371/journal.pcbi.1005530

Risse B, Thomas S, Otto N, Löpmeier T, Valkov D, Jiang X, Klämbt C. 2013. FIM, a Novel FTIR-Based Imaging Method for High Throughput Locomotion Analysis. PLoS One 8:1–11. doi:10.1371/journal.pone.0053963

Scialò F, Sriram A, Naudí A, Ayala V, Jové M, Pamplona R, Sanz A. 2015. Target of rapamycin activation predicts lifespan in fruit flies. Cell Cycle 14:2949–2958. doi:10.1080/15384101.2015.1071745

Sigrist SJ, Reiff DF, Thiel PR, Steinert JR, Schuster CM. 2003. Experience-Dependent Strengthening of Drosophila Neuromuscular Junctions. The Journal of Neuroscience 23:6546–6556. doi:10.1523/JNEUROSCI.23-16-06546.2003

Soiza-Reilly M, Azcurra JM. 2009. Developmental striatal critical period of activity-dependent plasticity is also a window of susceptibility for haloperidol induced adult motor alterations. Neurotoxicol Teratol 31:191–197. doi:10.1016/j.ntt.2009.03.001

Streit AK, Fan YN, Masullo L, Baines RA. 2016. Calcium Imaging of Neuronal Activity in Drosophila Can Identify Anticonvulsive Compounds. PLoS One 11:e0148461. doi: 10.1371/journal.pone.0148461

Sulkowski MJ, Han TH, Ott C, Wang Q, Verheyen EM, Lippincott-Schwartz J, Serpe M. 2016. A Novel, Noncanonical BMP Pathway Modulates Synapse Maturation at the Drosophila Neuromuscular Junction. PLoS Genet 12:e1005810. doi:10.1371/journal.pgen.1005810

Valdes-Aleman J, Fetter RD, Sales EC, Heckman EL, Venkatasubramanian L, Doe CQ, Landgraf M, Cardona A, Zlatic M. 2021. Comparative Connectomics Reveals How Partner Identity, Location, and Activity Specify Synaptic Connectivity in Drosophila. Neuron 109:105–122.e7. doi:10.1016/j.neuron.2020.10.004

Williamson M, Mitchell A, Condron B. 2021. Birth temperature followed by a visual critical period determines cooperative group membership. J Comp Physiol A Neuroethol Sens Neural Behav Physiol 207:739–746. doi:10.1007/s00359-021-01512-3

Winding M, Pedigo BD, Barnes CL, Patsolic HG, Park Y, Kazimiers T, Fushiki A, Andrade I V, Khandelwal A, Valdes-aleman J, Li F, Randel N, Barsotti E, Correia A, Fetter RD, Hartenstein V, Priebe CE, Vogelstein JT, Cardona A, Zlatic M. 2023. The connectome of an insect brain. Science (1979) 379. doi:10.1126/science.add9330

Zeng X, Komanome Y, Kawasaki T, Inada K, Jonaitis J, Pulver SR, Kazama H, Nose A. 2021. An electrically coupled pioneer circuit enables motor development via proprioceptive feedback in Drosophila embryos. Current Biology 31:5327–5340.e5. doi:10.1016/j.cub.2021.10.005

Zhong Y, Wu C-F. 2004. Neuronal Activity and Adenylyl Cyclase in Environment-Dependent Plasticity of Axonal Outgrowth in Drosophila. The Journal of Neuroscience 24:1439–1445. doi:10.1523/JNEUROSCI.0740-02.2004

Züfle P, Batista LL, Brandão SC, D’Uva G, Daniel C, Martelli C. 2025. Impact of developmental temperature on neural growth, connectivity, and function. Sci Adv 11. doi:10.1126/sciadv.adp9587

